# Transposable element landscape in *Drosophila* populations selected for longevity

**DOI:** 10.1101/867838

**Authors:** Daniel K. Fabian, Handan Melike Dönertaş, Matías Fuentealba, Linda Partridge, Janet M. Thornton

## Abstract

Transposable elements (TEs) inflict numerous negative effects on health and fitness as they replicate by integrating into new regions of the host genome. Even though organisms employ powerful mechanisms to demobilize TEs, transposons gradually lose repression during aging. The rising TE activity causes genomic instability and was implicated in age-dependent neurodegenerative diseases, inflammation and the determination of lifespan. It is therefore conceivable that long-lived individuals have improved TE silencing mechanisms resulting in reduced TE expression relative to their shorter-lived counterparts and fewer genomic insertions. Here, we test this hypothesis by performing the first genome-wide analysis of TE insertions and expression in populations of *Drosophila melanogaster* selected for longevity through late-life reproduction for 50-170 generations from four independent studies. Contrary to our expectation, TE families were generally more abundant in long-lived populations compared to non-selected controls. Although simulations showed that this was not expected under neutrality, we found little evidence for selection driving TE abundance differences. Additional RNA-seq analysis revealed a tendency for reducing TE expression in selected populations, which might be more important for lifespan than regulating genomic insertions. We further find limited evidence of parallel selection on genes related to TE regulation and transposition. However, telomeric TEs were genomically and transcriptionally more abundant in long-lived flies, suggesting improved telomere maintenance as a promising TE-mediated mechanism for prolonging lifespan. Our results provide a novel viewpoint indicating that reproduction at old age increases the opportunity of TEs to be passed on to the next generation with little impact on longevity.

## INTRODUCTION

Aging, also known as senescence, is an evolutionary conserved process described as the progressive loss of physiological homeostasis starting from maturity with disease promotion, decline in phenotypic function, and increased chance of mortality over time as a consequence (Fabian and Flatt 2011; Flatt and Heyland 2011; López-Otín et al. 2013). At the molecular level, studies of loss-of-function mutations in model organisms such as yeast, *Caenorhabditis elegans, Drosophila melanogaster*, and mice have successfully identified key pathways underlying aging and longevity including the conserved insulin/insulin-like growth factor signaling (IIS) and target of rapamycin (TOR) nutrient-sensing network (Piper et al. 2008; Fontana et al. 2010; Gems and Partridge 2013; Pan and Finkel 2017). More recently, sequencing of whole genomes, transcriptomes, and epigenomes corroborated that aging has a complex genetic basis involving many genes and is accompanied by changes across a broad range of interconnected molecular functions (López-Otín et al. 2013).

While there has been a predominant focus on understanding the links between genes and phenotypes correlated with aging, the role of transposable elements (TEs) in senescence and longevity has received less attention even though their discovery by Barbara McClintock goes back more than half a century ago (McClintock 1950). TEs, or transposons, are selfish genetic elements that replicate and move within genomes of their hosts. In eukaryotes, TEs typically constitute a considerable portion of the genome, with estimates around ∼3% in yeast, ∼20% in *D. melanogaster*, ∼70% in humans and ∼85% in maize (Quesneville et al. 2005; Schnable et al. 2009; de Koning et al. 2011; Carr et al. 2012). To date, several thousand TE families broadly classified into DNA-transposons and retrotransposons multiplying via RNA intermediates have been identified and are known to vary hugely in their transpositional mobility (Jurka et al. 2011; Deniz et al. 2019). For example, only a small fraction of L1 retrotransposons are responsible for most of the transposition events in the human genome, while the vast majority of L1s and other TE families have been inactivated by the accumulation of structural and point mutations over evolutionary time scales (Brouha et al. 2003).

In spite of the substantial evidence implicating TEs in adaptive evolution and diseases, the majority of transposons residing in the genome are likely to be neutral or only slightly deleterious for host fitness (Barrón et al. 2014; Arkhipova 2018). Yet, their exact physiological functions and the extent to which particular TE insertions or whole TE classes contribute to host fitness is still under debate (Brunet and Doolittle 2015). In general, TE mobility causes genomic instability through insertional mutagenesis, which can directly affect coding sequences of genes or modify their transcription. Typically, TE insertions into or close to genes impose negative consequences on health and have been associated with ∼100 diseases in humans, including cystic fibrosis, haemophilia and cancer (Hancks and Kazazian 2012). It is not just through the insertion of TEs that their presence may be deleterious, but also by causing detrimental chromosomal rearrangements resulting from ectopic recombination between TE families with similar sequences in different genomic locations (Montgomery et al. 1987; Charlesworth et al. 1992; Petrov et al. 2011). Additionally, TE expression and translation also allow the formation of toxic TE products that, for example, contribute to autoimmune diseases, while TE activity and replication of an increased genomic TE content might indirectly impose metabolic costs to the host (Kaneko et al. 2011; Barrón et al. 2014; Volkman and Stetson 2014; Bogu et al. 2019). On the other hand, there is mounting experimental evidence for positive selection on segregating TE insertions from multiple taxa confirming beneficial phenotypic properties including insecticide and virus resistance in *Drosophila* (Daborn et al. 2002; Magwire et al. 2011; Kuhn et al. 2014; Li et al. 2018; Rech et al. 2019).

A common feature of TEs observed in various organisms including yeast, *D. melanogaster, C. elegans*, mice, and humans, is the age-associated increase in transposition and expression, which usually coincides with weakening of the host TE silencing machinery and loss of genomic stability (Maxwell et al. 2011; Dennis et al. 2012; Solyom et al. 2012; De Cecco et al. 2013; Li et al. 2013; Gorbunova et al. 2014; Chen et al. 2016; Bogu et al. 2019; De Cecco et al. 2019). TEs have further been implicated in age-related neurodegenerative diseases (e.g. Krug et al. 2017; Prudencio et al. 2017; Guo et al. 2018) and might promote chronic inflammation observed during aging (Chen et al. 2014; De Cecco et al. 2019) further supporting the involvement of TEs in senescence and longevity as proposed by the emerging ‘transposable element theory of aging’ (Kirkwood 1989; Sedivy et al. 2013). The age-related change in TE activity detected in many tissues has mainly been attributed to chromatin remodeling and the decline in repressive heterochromatin structure which is commonly rich in transposable elements (Dimitri and Junakovic 1999; Wood and Helfand 2013; Chen et al. 2016; Wood et al. 2016). TEs that are not suppressed by chromatin structure are the target of post-transcriptional silencing by the host RNA-interference (RNAi) machinery, mostly the piwi-interacting RNA (piRNA) pathway, which is in turn also necessary for heterochromatin formation and stability (Lippman and Martienssen 2004; Martienssen and Moazed 2015). Indeed, research has identified longevity-promoting effects of several genes involved in the RNAi machinery and heterochromatin formation (Mori et al. 2012; Wood and Helfand 2013; Wood et al. 2016). Interestingly, it is possible that age-related misexpression of TEs is exclusive to the soma due to efficient post-transcriptional TE silencing mediated by the piRNA machinery in the germline (Sturm et al. 2015; Elsner et al. 2018; Erwin and Blumenstiel 2019). Considering current evidence, it seems natural that longevity can be achieved through impeding TE activity and controlling the genomic content of TEs. However, whether variation in aging and lifespan within species is also mediated by transposons and their role in the evolution of senescence is largely unknown.

Here, we analyze genomes of *D. melanogaster* populations experimentally selected for increased lifespan through postponed reproduction from four independent studies to understand the role of TEs in the evolution and genomic basis of late-life performance and aging. The invertebrate *D. melanogaster* is an excellent model in this respect as it exhibits abundant genetic and phenotypic variation in fecundity and traits related to aging that can be selected for. In the present experiments, replicate populations derived from nature were subjected to a late-life breeding scheme in which only flies surviving and fertile at old age contributed to the subsequent generations, while control individuals reproduced earlier in life. When the genomes were sequenced, the selection process had continued for over 30 years with ∼170 and ∼150 generations of selection for Carnes et al. 2015 (Carnes2015) and Fabian et al. 2018 (Fabian2018), and for 58 and 50 generations for Hoedjes et al. 2019 (Hoedjes2019) and Remolina et al. 2012 (Remolina2012) enabling us to quantify differences in TE content of long- and short-term evolutionary responses. Selection for postponed senescence has resulted in phenotypic divergence of multiple fitness traits, most notably an ∼8% to ∼74% increase in lifespan and improved old age fecundity at the cost of reduced early reproduction (Luckinbill et al. 1984; Rose 1984; Remolina et al. 2012; Carnes et al. 2015; Fabian et al. 2018; Hoedjes et al. 2019; May et al. 2019). At the genome level, analysis of genetic differentiation has revealed a significant sharing in candidate genes across the four studies indicating parallel evolution (Hoedjes et al. 2019), but at the same time exposed multiple novel targets of selection. For instance, three of the studies report genetic and/or transcriptomic divergence in immunity genes, and it has recently been confirmed that these molecular changes reflect differences in traits related to pathogen resistance (Fabian et al. 2018). Thus, despite variations in the experimental designs, numerous evolutionary repeatable phenotypic and genetic adaptations have been observed, but the importance of TEs in these studies has remained unexplored. Therefore, our main objective was to investigate for the first time whether TE abundance in the genome, and host genes related to TE regulation, had undergone similar parallel changes. Using RNA-seq data from Carnes et al. (2015), we further test if males and females of selected populations evolved to suppress TE transcription to mitigate potentially negative effects on longevity.

## RESULTS

### Selection for postponed reproduction affects genomic abundance of TE families

To analyze if selection for longevity affected TE copy number, we used DeviaTE (Weilguny and Kofler 2019) on whole genome pool-sequences of a total of 24 late-breeding, long-lived selection (S) and 22 early-breeding control (C) populations from four studies (see **Table S1** for details on experimental designs) (Remolina et al. 2012; Carnes et al. 2015; Fabian et al. 2018; Hoedjes et al. 2019). DeviaTE is an assembly-free tool that estimates genomic abundance of 179 TE families by contrasting the sequencing depth of TEs and five single-copy genes taking internal deletions within TEs into account (**Fig. S1** and **Fig. S2**).

After employing coverage and mapping quality filters (**Fig. S3**), we screened for differences in abundance between control and selection regimes of 110 to 115 TE families dependent on the study, using three complementary approaches that vary in stringency (see overview in **Fig. S1** and Materials and Methods, summary statistics in **Table S2**). In brief, we (1) analyzed studies independently, (2) fit models combining all studies using proportions of TE family abundance relative to the total genomic TE content, and (3) tested if copy number differences are driven by TE expansions specific to particular populations by investigating if changes in TE abundance are consistent across all replicates within regime and study. For all methods, we found more TE families with higher copy numbers in selected populations relative to controls than vice versa, with the exception of the high protein/sugar larval diet regime in Hoedjes2019 (**Table 1**, see **Supplementary Results**, for breeding regime differences within each diet also see **Table S3** and **Fig. S4**). We further obtained qualitatively similar results when we only considered the last 200 bp at the 3’-ends of the TE families, which are thought to harbor less deletions and truncations (**Table S4**), and when we analyzed sequence abundance using sums of normalized coverage values across the TE family consensus (**Table S5**).

**Table 1.**
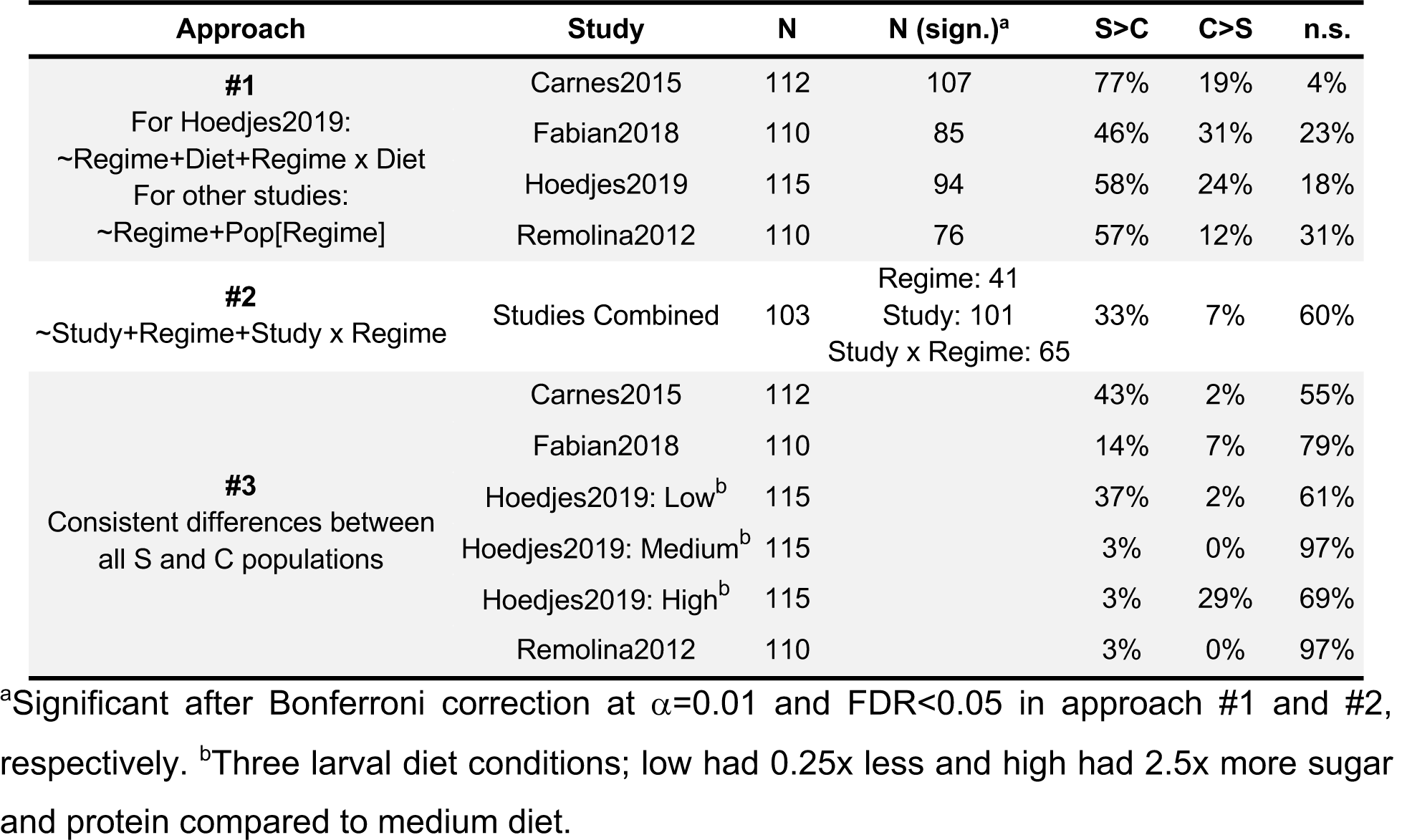
Number of detected TE families (N) and percentage of families more abundant in selected (S>C) or control regimes (C>S) or not different (n.s.) using three different approaches (also see Fig. S1 and Table S2).

For the downstream analysis, we describe TE families varying between regimes as defined by approach #1 (**Fig. 1A, Table 1**). In this approach, between 46% and 77% of all TEs had a significantly larger number of genomic insertions in the selected populations relative to controls after Bonferroni correction for multiple testing (from here on referred to as S>C TEs). In contrast, only 12% - 31% of TEs showed the opposite pattern and had more insertions in the controls (from here on referred to as C>S TEs).

**Figure 1.**
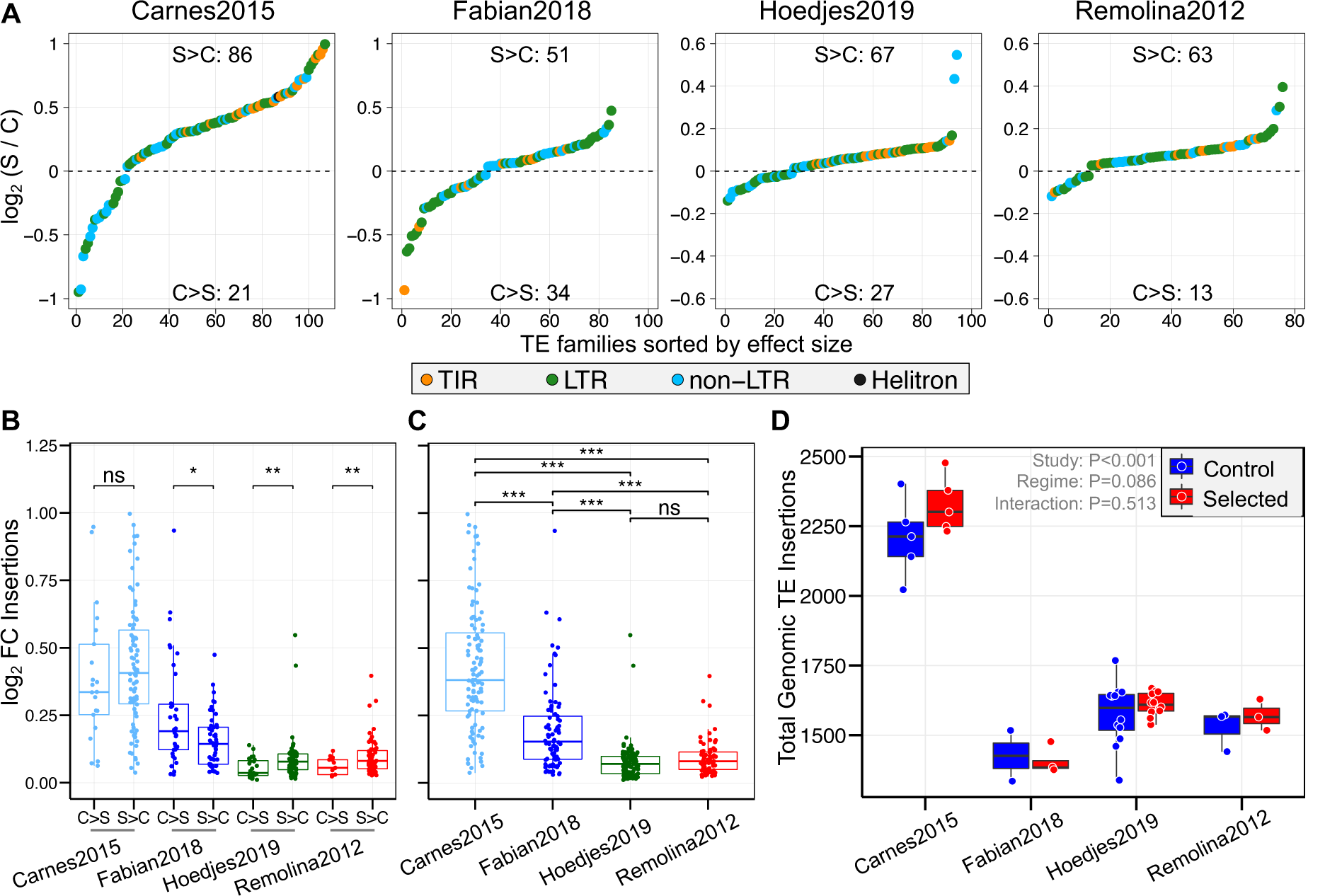
Dynamics of TE copy number change between breeding regimes. (A) Log_2_ fold change in average genomic insertions of the late-breeding selected populations (“S”) relative to early-breeding controls (“C”). The dashed line indicates no difference between regimes. >0 denote TE families with a larger abundance in selected populations (“S>C”), while <0 TEs with more insertions in controls (“C>S”). Number of TE families in these two categories are given in the center at the top and bottom of each plot. TE subclasses are given in different colors. Selected flies had more genomic insertions than controls for most TE families (also see **Table 1**). (B) Difference in the magnitude of absolute log_2_ fold change between C>S and S>C TE groups. Significant difference between TE groups was determined using t-tests for each study. (C) Magnitude of absolute log_2_ fold change between studies, analyzed using ANOVA with Study as single term (*F*_3,358_ = 106.5, P < 2e-16) and pairwise Tukey post-hoc tests. * P < 0.05; ** P < 0.01; *** P < 0.001; ns, not significant. (D) Total number of genomic TE insertions. We used ANOVA to test the effects of Study, Regime and the Study x Regime interaction. See **Table S6** for a summary of the statistical analysis.

To explore if the dynamics of TE copy number change are similar among studies, we first contrasted log_2_ fold changes in abundance between S>C and C>S TEs. S>C TEs had a significantly larger magnitude of change than C>S TEs in the two short-term evolution studies of Hoedjes2019 and Remolina2012, while the opposite pattern was observed for Fabian2018 and no difference for Carnes2015 (**Fig. 1B**; t-tests, all P < 0.05 except Carnes2015: P = 0.466). Moreover, studies differed significantly in the size of log_2_ FC values in the order of Carnes2015 > Fabian2018 > Hoedjes2019 = Remolina2012 (**Fig. 1C**; ANOVA with Study term, Tukey HSD, P < 0.001 for all pairwise comparisons except Hoedjes2019-Remolina2012, P = 0.924), seemingly scaling with the length of selection (Carnes2015: 170; Fabian2018: ∼146; Hoedjes2019: 58, Remolina2012: 50 generations).

We next asked if changes in TE abundance are driven by certain TE subclasses (Long Terminal Repeat, LTR; Non-Long Terminal Repeat, non-LTR; Terminal Inverted Repeat, TIR) or class (RNA, DNA) and tested S>C and C>S TEs for enrichment of these types using two-sided Fisher’s exact tests. We only detected a significant underrepresentation of TIRs and DNA-class TEs (i.e. overrepresentation of RNA-class) in the C>S group of TEs of Carnes2015 and Hoedjes2019 (Carnes2015, TIRs: P = 0.044; DNA/RNA class: P = 0.024; and Hoedjes2019, TIRs: P = 0.013; DNA/RNA class: P = 0.008), while there was no enrichment in Fabian2018 and Remolina2015.

Despite many individual TEs having a higher genomic abundance in the selected populations, the whole genomic TE content was not significantly different between the regimes, but varied among studies (**Fig. 1D**, and **Table S6**). This was at least partly driven by the fact that although C>S TEs were fewer in number than S>C TEs, they showed a significantly higher difference in insertion counts in two studies (**Fig. S5**, t-test using *δ*Insertion values; Carnes2015 P = 0.04; Fabian2018 P = 0.005). The non-significant difference in overall genomic TE load could therefore be a result of a large number of S>C TEs with small differences that are balanced by fewer C>S TEs with large differences. We further analyzed the whole genomic abundance of individual subclasses of TEs and identified a significantly higher TIR content in selected populations compared to controls (**Fig. S6**, ANOVA, both Regime and Regime x Study factors, P < 0.001), but this effect was strongly influenced by Carnes2015 (Tukey HSD, Regime x Study factor testing for C vs S within studies, Carnes2015: P < 0.0001; other studies: P > 0.85). We also detected that selected populations had a larger LTR retrotransposon load than controls (ANOVA, Regime factor, P = 0.026), whereas non-LTR content did not differ significantly. Finally, we note that studies in general varied significantly in total TE content and subclass-specific loads (ANOVA, Study factor, P < 0.0001 in all models).

In summary, our results demonstrate that selection for postponed reproduction leads to evolutionary repeatable increases in copy number of many TE families relative to early bred controls, but without affecting the overall genomic TE load.

### TE families varying in genomic abundance differ in evolutionary age and activity

We next tested if differences in TE activity explain the changes in abundance between control and selected populations. In *Drosophila*, most TE families are considered to be active (Guio and González 2019), and it has been shown that the average population frequency of TE insertions within a family serves as a good proxy for recent activity and age of TE invasion (Kofler et al. 2012; Kofler et al. 2015).

We first determined the exact genomic location and frequency of TE insertions using PoPoolationTE2 (Kofler et al. 2016) and calculated average population frequency across all insertion sites for each TE family. As expected, the number of detected TE insertions which could be mapped to genomic locations partially scaled with coverage (see Materials and Methods): across all populations within a study, we found 13,018 TE insertions in Hoedjes2019, 8,402 in Fabian2018, and 4,502 in Remolina2012, which is in the range recently identified in natural populations (i.e. 4,277 - 11,649 TE insertions in Lerat et al. 2019). The least number of TE insertion locations was found for Carnes2015 for which we detected an unusually small number of 567 TE insertions, likely reflecting a large number of false negatives due to low sequencing depth. For each TE family, we then averaged frequencies across all of its detected genomic positions to estimate the mean frequency at which a TE is segregating in a population (Kofler et al. 2015). Studies varied in the minimum average TE family frequency in the order of Carnes2015 > Remolina2012 > Fabian2018 > Hoedjes2019, which is likely a further effect of dissimilar sequencing depths and other experimental factors (average frequency ranges of Hoedjes2019: 0.01 - 0.9; Fabian2018: 0.02 - 1; Remolina2012: 0.04 - 0.84; Carnes2015: 0.19 – 0.9). Therefore, the TE frequencies of Carnes2015 need to be interpreted with care, considering the likely insufficient amount of data.

To get unbiased average TE frequency estimates independent of coverage fluctuations across studies, we also obtained average frequencies from a single natural South African (SA) population (Kofler et al. 2015; Kofler 2019). The SA population had a higher sequencing depth than all studies here (i.e. 381x) and thus presumably a more accurate estimate of TE frequencies. Notably, this population was not subjected to any selection or control treatment and was only maintained 8 generations in the lab before sequencing. Average genome-wide TE frequencies of control and selected populations of Fabian2018, Hoedjes2019 and Remolina2012, but not Carnes2015, were significantly correlated with the South African TE frequencies (**Fig. S7**; Spearman’s ρ, Fabian2018: 0.65; Hoedjes2019: 0.61; Remolina2012: 0.58, all three P < 0.0001; Carnes2015: 0.1, P = 0.403), demonstrating that the SA population can function as an appropriate reference here.

In accordance with previous reports, we confirmed that the TE content of all populations consists of a large number of low frequency and fewer high frequency TE families (**Fig. S8**, Spearman’s ρ between TE abundance and average frequency of SA population, ρ = -0.4 to -0.54, all P < 0.0001; similar when frequencies of experimental evolution studies were used, see **Fig. S9**) (Petrov et al. 2011; Kofler et al. 2015).

We then examined the data of the SA population and found that C>S TEs had a significantly lower frequency than S>C TEs in all four studies (**Fig. 2**, t-tests between C>S and S>C frequencies, P < 0.05 for all four studies). As there were more S>C than C>S TEs, we also contrasted the average frequencies of the top 10 C>S and S>C TEs with the biggest changes in genomic abundance defined by log_2_ FC values (**Fig. 1A**). We only detected a significantly higher frequency in top 10 S>C relative to C>S TEs for Carnes2015 (t-test, P = 0.03), but not in the other three studies. Considering the relationship between insertion age, frequency and activity of TE families (Kofler et al. 2015), the lower frequency of C>S TEs suggests that they are evolutionary younger and potentially more active than S>C TEs.

**Figure 2.**
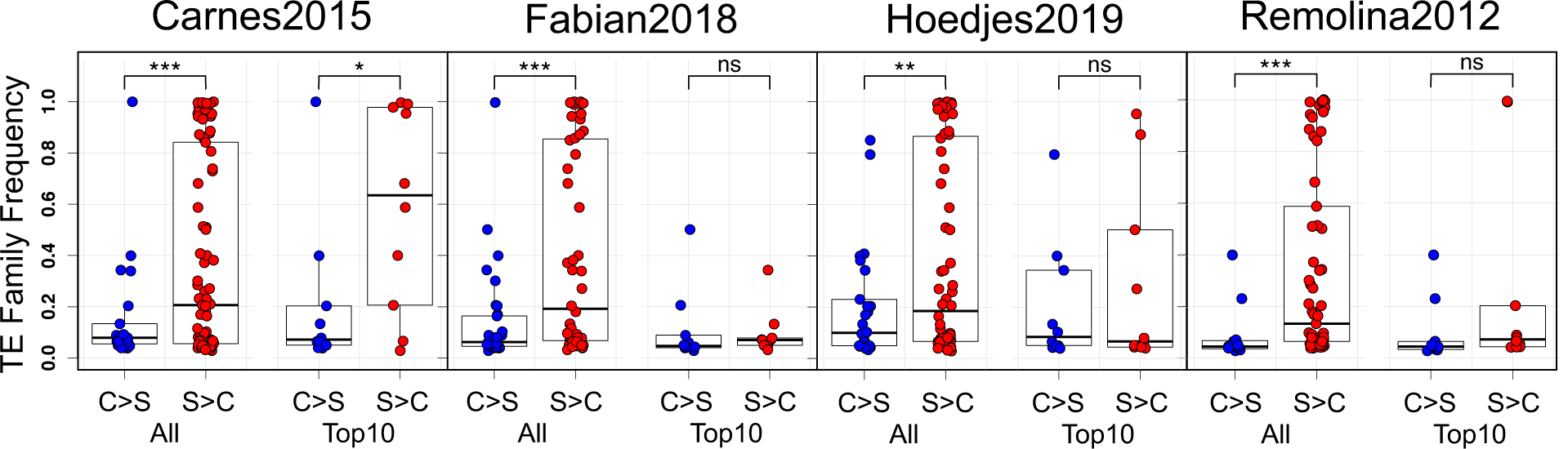
Differences in average TE frequency. Average TE frequency from the South African population separated into C>S (blue) and S>C TEs (red) are shown on the Y-axis. We investigated differences considering all C>S and S>C TEs (“All”) or only the top 10 TEs with the biggest differences in log_2_ FC of insertions (“Top 10”). t-tests were used to assess statistical significance. ns, not significant; * P < 0.05; ** P < 0.01; *** P < 0.001.

### Genetic drift is not driving differences in TE abundance

A major challenge in experimental evolution studies is to differentiate selection from the confounding genomic signals of genetic drift, which might be amplified by small effective population sizes (*N*_*e*_) or varying generation times spent in the lab between control and selected populations. We therefore calculated genome-wide nucleotide diversity *π* and Watterson’s *θ* across 100kb windows as a proxy for *N*_*e*_. With the exception of Fabian2018, where *π* was equal between regimes (ANOVA, Regime factor, P = 0.179), we found that both estimators were significantly higher in selected relative to control populations (**Table S7**; ANOVA, Regime factor, all P < 0.0001). Even though a generally reduced *N*_*e*_ in controls should lead to the loss of low and fixation of high frequency TEs under neutrality, we observed the opposite pattern in our analysis above (**Fig. 2**).

To further formally test if the increased abundance of many TEs is driven by selection on preexisting TE insertions or genetic drift alone, we performed population genetic simulations using the correlated average TE frequencies from the natural South African population (Kofler et al. 2015) as a starting point (see **Fig. S7** and results above). We simulated TE frequency change in selected and control populations 5,000 times given the reported consensus population sizes as *N*_*e*_, generation times and number of replicates. We then asked how often the same or a higher relative proportion of S>C to C>S TEs as in our observations is obtained (**Table 1**). While the results from Carnes2015, Hoedjes2019, and Remolina2012 were significantly different from the expected proportions, the TE abundance differences of Fabian2018 could be caused by genetic drift alone (**Fig. S10**). Testing different ranges of the reported population sizes and assuming that only 50% and 25% of flies in the selected populations were able to breed at old age resulted in qualitatively similar results (not shown). We also quantified expected proportions of TEs consistently varying in frequency across simulated replicates: while there were generally more TEs consistently higher in abundance in selected populations (**Table 1**, approach #3), all our simulations resulted in more TE families with a consistently higher frequency in controls. The increased genomic abundance of many TEs in selected populations is therefore unlikely to be solely caused by genetic drift.

### Limited evidence for selection on TE abundance and insertion frequencies

Considering the deviation from neutrality, we next asked if the parallel patterns in TE abundance are caused by the same or different TEs, which could indicate selection acting on genomic copy number of certain TEs. Among the 103 common TE families, we identified 14 S>C and 2 C>S TEs shared across all four studies (**Fig. 3A, Table S8**). Despite the seemingly large number of shared S>C TEs, only the overlap between Remolina2012 and Hoedjes2019 was significant (P = 0.025). Yet, we found that the most common telomeric TE *HeT-A* (Casacuberta 2017) was on average more abundant in selected populations in all four studies (**Fig. S1**, also identified by approach #2, see **Table S2**), suggesting that long-lived populations might have evolved longer telomeres to avoid attrition, which is considered to be a key conserved mechanism of aging (López-Otín et al. 2013). In contrast to S>C TEs, C>S TEs showed significant overlaps across all four studies, two triple set comparisons, and between Remolina2012 and Hoedjes2019 (**Fig. 3A, Table S2**). Potentially, a high genomic abundance of *G-element* and *G2* found in the control populations of all studies is detrimental for longevity and late-reproduction (**Fig. 3B**). However, we did not observe any significant Spearman’s correlation coefficients in pairwise comparisons of log_2_ FC values between studies except for Hoedjes2019-Remolina2012 (ρ = 0.28, P = 0.004), showing that TE families generally lack parallel changes in abundance.

**Figure 3.**
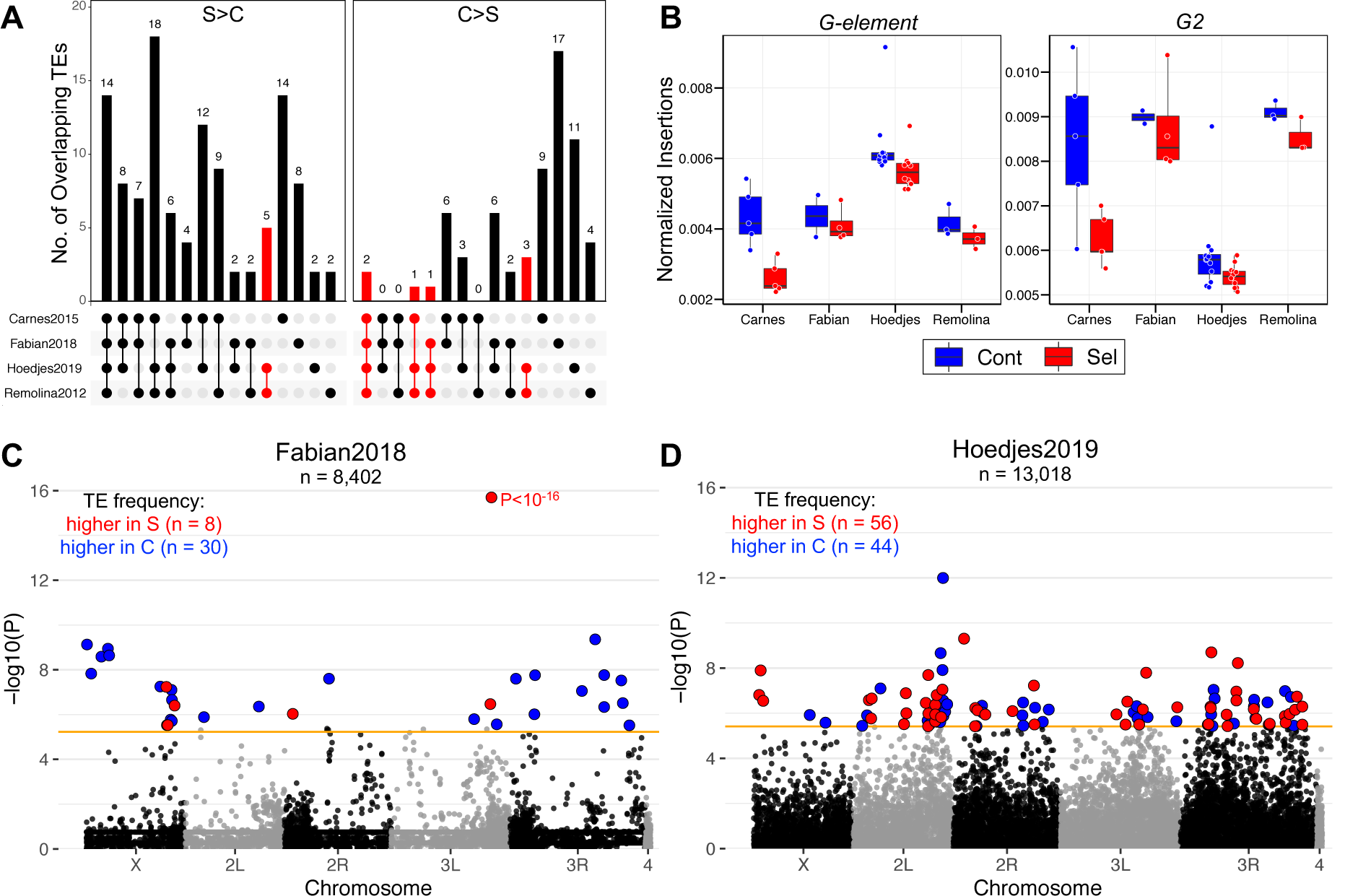
Selection on TE abundance and insertions. (A) Overlap of TE families with significant abundance differences among studies. S>C and C>S denote TEs with a higher abundance in selected or control populations, respectively. Red bars indicate a significant overlap at P < 0.05 (also see **Table S8**). (B) Boxplots of the number of genomic insertions relative to the total genomic content of the 2 significantly shared C>S TEs. (C) Genome-wide differentiation in TE insertion frequency between selected and control populations in Fabian2018 and (D) Hoedjes2019. Every point indicates the -log_10_ P-value of a TE insertion across chromosomal arms (alternating black and grey color). The solid orange line corresponds to the Bonferroni cut-off at *α* = 0.05 (Fabian2018: P < 5.9 × 10^−6^; Hoedjes2019: P < 3.8 × 10^−6^). Red and blue points denote TE insertions with a significantly higher frequency in selected or control populations, respectively. More details including exact positions, frequency and annotation of candidate TE insertions can be found in **Table S9**.

Genomic TE abundance in selected populations might also be increased because selection acted on a large number of segregating TE insertions resulting in frequency divergence between control and selected populations. We therefore screened all identified TE insertion sites for significant frequency differences between regimes in each study by performing ANOVAs on arcsine square root transformed frequencies **(Table S9**). After correcting for multiple testing, we detected significant frequency differences for 38 TE insertions in Fabian2018 and 100 in Hoedjes2019 (**Fig. 3C** and **Fig. 3D**). At the gene level, the significant TEs defined 29 and 98 genes in Fabian2018 and Hoedjes2019, respectively, and none were shared between the two studies. However, in Carnes2015 and Remolina2012 insertions did not show significant frequency differentiation even at a less stringent cut-off (FDR < 0.05). We further tested if TE families varying in genomic abundance also differ significantly in frequency between the regimes (**Table S10**, see **Table S11** for statistics on each TE family). There was little evidence for parallel patterns in all studies except from Carnes2015 (Carnes2015: 27 TE families significant for abundance and frequency; other studies: 0 to 3). Thus, although differences in TE abundance are unlikely to be driven by neutral evolution alone, we only found limited evidence for parallel evolution of TE copy numbers and sparse TE frequency differentiation.

### Sex, age, and selection regime affect TE expression

To test whether the increased genomic abundance of TE families in selected flies is explained by a higher transcriptional activity we analyzed RNA-seq data from whole flies of Carnes2015 (**Fig. 4** and **Table S12**, see **Table S13** for the complete statistical analysis). We first fit a model with Sex, Age, and Regime to every TE family and each gene on the major chromosomal arms **(Fig. S11)**. In line with sex differences in gene expression observed by Carnes et al. (2015), ∼92% of TE families had a significant sex term of which most had a higher expression in males than females.

**Figure 4.**
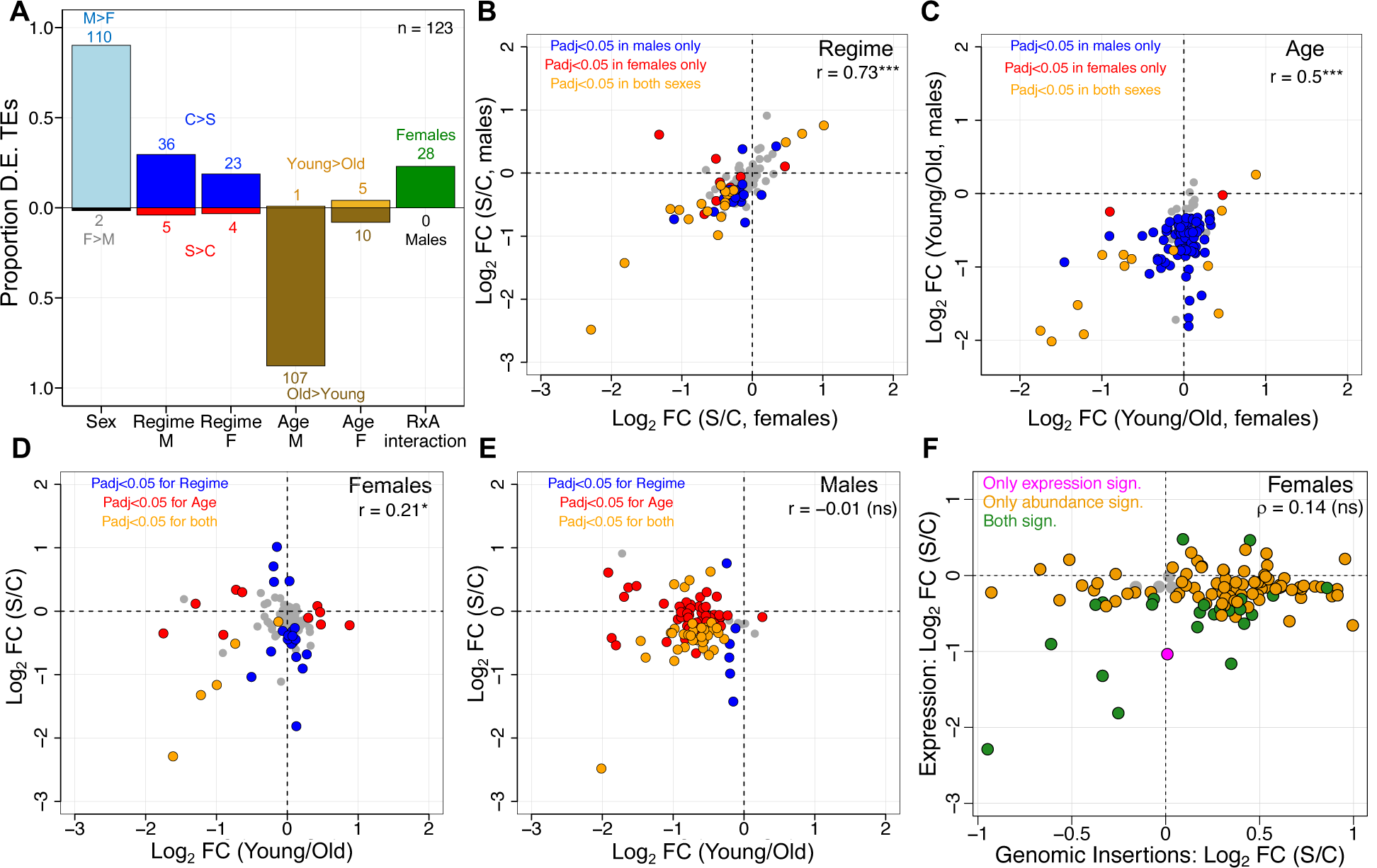
Multiple factors influence TE expression. (A) Proportions of differentially expressed TEs at adjusted P < 0.05 and directionality relative to 123 TEs with detectable expression for factors from statistical models on pre-filtered read counts in DESeq2 (also see **Table S12**). “Sex” refers to the results of the model including Sex (M, males; F, females), Age (young; old), and Regime (C, control; S, selected). “Regime”, “Age” and “RxA” (i.e. Regime x Age interaction) refer to results from model fits with males and females separately analyzed. The absolute number of TEs for factor levels are given above or below bars. (B) Log_2_ fold change of regime (selected vs control) and (C) age (young vs old) for males and females. Colors designate TEs significant only in males (blue), or females (red), or shared between both sexes (orange). Not significant TEs are in grey. (D and E) Log_2_ fold changes across regime against age differences in males and females. Colors designate TEs significant only for regime (blue), or age (red), or for both factors (orange). Not significant TEs are in grey. (F) Relationship of log_2_ fold changes in TE expression and genomic abundance between regimes in females. (B to E): r, Pearson’s correlation coefficient; (F): ρ, Spearman’s correlation coefficient; * P < 0.05; *** P < 0.0001; ns, not significant.

We therefore decided to test the effects of Regime, Age, and the Regime x Age interaction in the sexes separately (**Fig. 4A, Table S12**). We detected 41 (∼34% of total) and 27 TEs (∼22%) significantly different between regimes in males and females, respectively, with the majority being upregulated in controls (**Fig. 4B**). Among these, 19 TEs significant in both sexes also had the same directionality of expression change: 10 LTR-class TEs and 6 non-LTRs were higher expressed in controls, whereas 3 non-LTR TEs (*TART-A, TART-B, and TAHRE*) were upregulated in selected populations **(Table S14)**. Interestingly, *TART-A, TART-B*, and *TAHRE* provide the enzymatic machinery for telomeric maintenance (Casacuberta 2017), again suggesting that reduced telomere attrition evolved in response to selection, paralleling the genome-based analysis. In general, regime affected TE expression in males and females similarly, as indicated by a significant correlation of log_2_ fold change values between sexes (**Fig. 4B**, Pearson’s r = 0.73, P < 0.0001). We further asked if the magnitude of log_2_ fold change varies between TEs more expressed in controls or selected populations, and did not find any significant difference (**Fig. S12**, t-test, females: P = 0.86; males: P = 0.95).

Supporting the notion that TEs become derepressed during aging, the effect of age on TE expression in males was general as 107 of the 108 significant TEs (i.e. ∼88% of all included TE families) had a higher expression in older flies. Less pronounced differences were found in females where 8% of all TEs – all of which were retrotransposons – increased and 4% of TEs decreased expression with age (**Fig. 4A** and **Fig. 4C**). Moreover, consistent with a recent study (Chen et al. 2016), the TEs upregulated in older females had on average a significantly higher log_2_ fold change relative to the downregulated TEs (**Fig. S12**, t-test, P = 0.018). We further found 13 TEs with a significant age factor in both sexes (**Fig. 4C, Table S15**), of which *copia, Burdock, R1* and *R2* are already known to increase expression with age (Li et al. 2013; Chen et al. 2016).

No TE families showed a significant Regime x Age term in males, but the interaction was significant for 28 TEs (∼23% of total) in females (**Fig. 4A**). Interestingly, most of these TEs were defined by a higher expression in young controls compared with selected flies of the same age (see **Fig. S13** for example). Selected populations subsequently increased while controls decreased expression, meeting at a similar expression level at old age. This is comparable with recent studies which suggested that age-dependent changes in TE expression differ between genotypes (Erwin and Blumenstiel 2019; Everett et al. 2020).

We next investigated if differential expression of TEs is specific or similar to the overall transcriptomic changes by comparing proportions of TEs and genes up- or downregulated or unchanged within levels of sex, regime, and age (**Fig. S14**). Distributions generally varied significantly (*χ*^2^ tests, P < 0.001 for all, except age factor in females: P = 0.129), demonstrating that these factors have different effects on TE and gene expression.

To further examine if the selected populations might have evolved to maintain a young TE expression profile, we compared differences between regimes to those that occurred with age (**Fig. 4D and 4E**). The correlation of log_2_ FC values between regime and age was positive for TEs in females (Pearson’s correlation, females: r = 0.21, P = 0.021; males: r = -0.01, P = 0.875), and varied from the one for genes (1000 bootstrap replicates resampling 100 genes: mean Pearson’s correlation, females: r = -0.12, 95% CI: -0.13 to -0.11; males: r = 0.09, 95% CI: 0.08 to 0.1). Thus, expression of TEs between selected and control populations only mirrors the changes between young and old flies in females.

In summary, our results suggest that selected populations of Carnes2015 evolved to reduce TE expression, but differences across sex and age were overall more dominant than variation between regimes.

### Differences in TE abundance do not match TE expression patterns

We also asked if the change in genomic TE abundance parallels the expression differences between selected and control populations. Notably, as the genomic TE abundance measures came from DNA pools of female flies, we did not do this comparison in males. We first confirmed that TE expression scaled robustly with the number of genomic insertions in each age-regime combination (Spearman’s ρ = 0.72; P < 0.0001; **Fig. S15** and **Table S16**). Next, we investigated if there were parallel changes in 23 TEs significantly varying between regimes in expression and genomic abundance. We found that a majority of 13 TE families had non-parallel changes (**Table S17**). Indeed, log_2_ FC expression and log_2_ FC insertions between regimes were not significantly correlated (**Fig. 4F**, Spearman’s ρ = 0.14, P = 0.149), indicating that differences in TE abundance poorly predict differential expression between control and selected populations. As expected, correcting RNA-seq read counts for TE copy number to examine if average expression per TE insertion varies between regimes yielded qualitatively similar results compared to analyzing overall TE expression (**Table S18**). However, the tendency of TE families to be more highly expressed in controls was substantially larger (63 TEs more, 3 TEs less expressed in controls), further emphasizing that selection for late-reproduction leads to a reduction in TE expression.

### Little study-wide sharing in candidate genes involved in regulation of TE activity

We next hypothesized that if TE expression and transposition are predominantly detrimental for lifespan and aging, as proposed by many studies, experimental evolution for longevity would have likely resulted in selection on host alleles that influence TE activity. To test this, we screened 96 chromatin-structure, piRNA, and transposition-associated genes known to be involved in TE regulation and silencing for clear-cut genetic and expression differentiation possibly driven by selection (**Table S19**). Of these, 3 to 10 genes were implicated under selection across the four studies, and only *E2f1* (FBgn0011766) and *Hsp83* (FBgn0001233) were shared between two datasets (**Fig. 5A**). Moreover, the four studies did not report any significant enrichment of GO terms related to transposon silencing and chromatin structure. Using the available RNA-seq data from whole flies of both sexes in Carnes2015 and microarray data from female heads and abdomens in Remolina2012, we then asked if TE regulation genes are differentially expressed (**Fig. 5B**). We found that the 42 TE regulation genes significant for regime tended to be upregulated in controls in Carnes2015, but only two genes differed in Remolina2012. Interestingly, similar to TE expression patterns in Carnes2015 (**Fig. 4A**), TE regulation genes showed a clear tendency for upregulation with age in males but to a lesser degree in females (**Fig. 5B**). Comparable patterns were detected in Remolina2012, where the age effect was stronger in abdominal compared to head tissue. Thus, the boosted expression of TE regulation genes at older ages appears to be common and might be a response to increased TE transcription in old flies.

**Figure 5.**
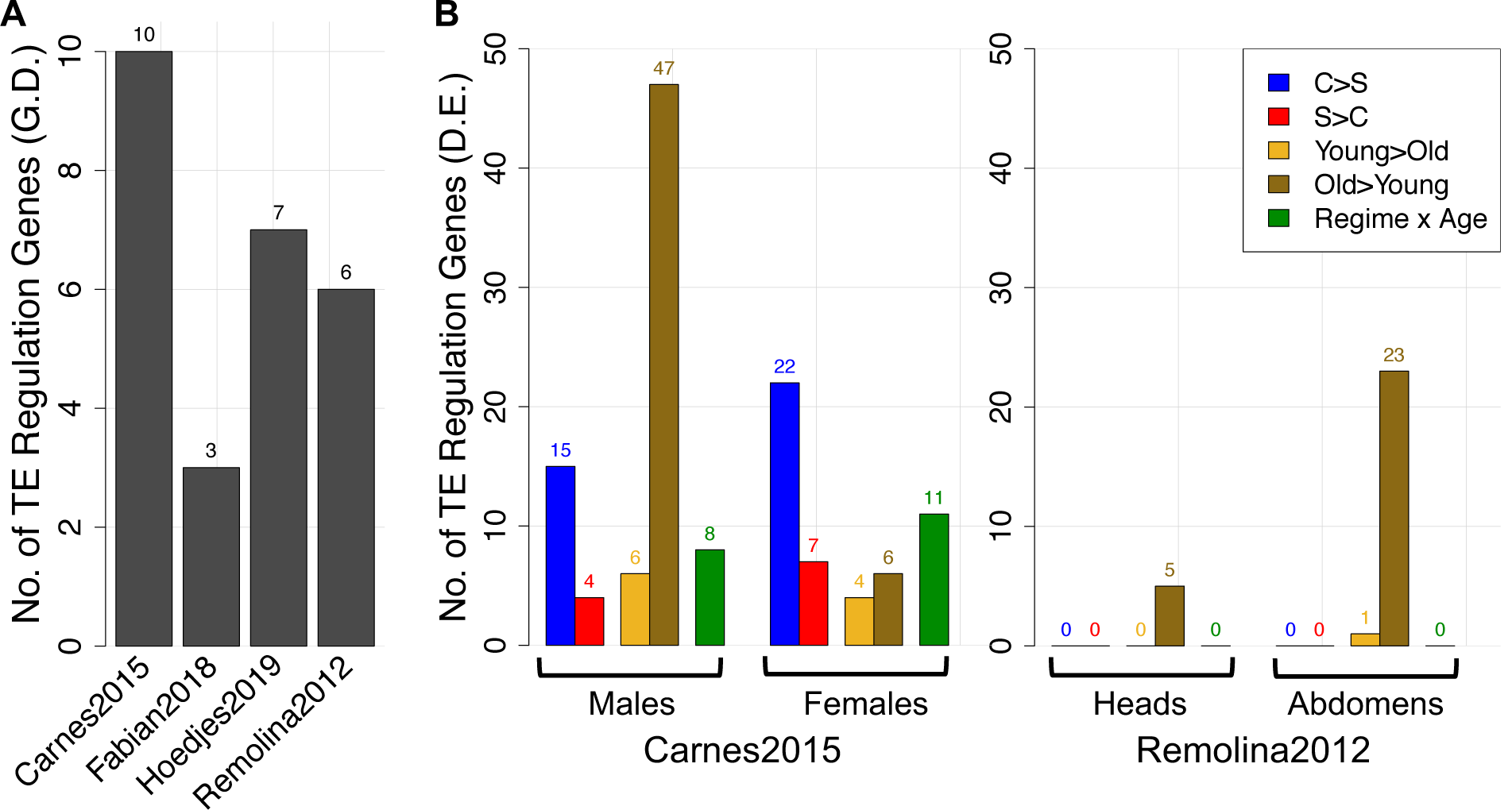
Number of genetically and transcriptionally differentiated genes involved in regulation of TE activity. (A) Counts of genetically differentiated (G.D.) TE regulation genes reported in the four experimental evolution studies. (B) Number of TE regulation genes differentially expressed (D.E.) between regimes (C, control; S, selected) and ages (young; old) in the RNA-seq data of Carnes2015 (whole female flies) and microarray data of Remolina2012 (female heads and abdomens). Also see **Table S19** for information on all 96 TE regulation genes.

Taken together, the small number of genetically differentiated TE regulation genes, lack of TE-associated GO enrichment, and overall missing parallel patterns suggest that improving TE repression was either specific to studies and/or not a prime target of selection.

## DISCUSSION

Are transposable elements conferring an adaptive advantage as shown for many traits (Daborn et al. 2002; Magwire et al. 2011; Kuhn et al. 2014; Li et al. 2018; Rech et al. 2019) or should they be purged and repressed during the evolution of longevity due to their widespread negative effects on fitness (Chen et al. 2014; Krug et al. 2017; Prudencio et al. 2017; Guo et al. 2018; De Cecco et al. 2019)? In this report, we attempt to answer this controversial question by employing four independent data sets to present the first characterization of the genome-wide TE content and expression in *D. melanogaster* populations that were experimentally selected for late-life reproduction and longevity.

### Does longevity-selection lead to changes in TE abundance?

Variation in TE copy number has been associated with some geographic and climatic factors (Kalendar et al. 2000; Kreiner and Wright 2018; Lerat et al. 2019) in natural populations of plants and *Drosophila* and was shown to change during experimental evolution in different temperatures (Kofler et al. 2018). Our analysis revealed a repeatable trend showing that many, but not all, TE families have an increased number of genomic insertions in late-breeding, long-lived populations, which indicates that reproductive age, with some dependency on developmental diet, is another factor influencing divergence in TE abundance **(Fig. 1A** and **Table 1)**. Interestingly, we found a significant difference in the magnitude of TE abundance change between studies that roughly scaled with the number of generations under selection (**Fig. 1C**). While parallel changes in TE characteristics within populations of the same selection regime have been reported by similar experiments (Graves et al. 2017; Kofler et al. 2018), it is striking that we observed this pattern in data created by four independent studies. Despite a lot of TE families being more abundant in long-lived populations, our analysis shows no significant difference in the total genomic TE content between control and selected populations (**Fig. 1D**), possibly because there were a few TEs with large increases in copy number in controls in contrast to many TEs with small increases in abundance in selected populations (**Fig. S5**). Changes in the overall genomic TE load are therefore likely not essential to evolve longevity or fecundity at old age in *Drosophila*. These findings are in contrast to recent work in several killifish species, which reported that TE expansion can cause an increased genome size with possible negative effects on lifespan (Cui et al. 2019). However, our analyses focused exclusively on the genomic TE load and as such we cannot exclude a difference in genome size between control and selected populations, which may be caused by other factors such as non-repetitive InDels or repetitive DNA unrelated to TEs.

### Are TEs adaptive during the evolution of aging?

The genomic content of TEs evolves through various factors, including replicative transposition, selection, genetic drift, and the TE defense machineries of the host (Charlesworth and Charlesworth 1983; Kofler 2019). By performing population genetic simulations that consider only genetic drift, we were able to exclude that population size and generations spent in the lab *per se* cause an increased abundance of TE families in selected populations (**Fig. S10**). Even though it is known that the majority of TE insertions are neutral to fitness (Arkhipova 2018), our findings suggest that factors other than genetic drift influenced TEs.

From a selective point of view, increasing many TE families might be beneficial for longevity, while fewer families could affect lifespan negatively. Under this scenario, selection would lead to parallel increases or decreases of the same TE families across studies. However, when we screened for parallel patterns in abundance change, we found only two TEs (*G-element* and *G2*) that had decreased copy numbers in selected flies and were significantly shared across all studies (**Fig. 3A,B)**. Both elements are *jockey*-like non-LTR TEs, of which *G2* is highly enriched in centromeric regions of the genome (Chang et al. 2019). Thus, changing centromeric structure by altering its TE content could be one mechanism modulating aging, although experimental evidence for this is still missing. In contrast to this, we did not find any significant overlap between all four studies among TEs with an increased abundance in the late-breeding populations. Unless many TE families had non-repeatable effects on longevity, the small amount of significant sharing suggests that abundance of most TEs is neutral.

Another possibility is that TE abundance is altered through selection affecting TE insertions at a genome-wide scale, resulting in a large number of insertions significantly varying in frequency between control and selected populations. We found only a minor fraction of TE insertions in Fabian2018 and Hoedjes2019, but not in the other two studies, with significantly different frequencies between the regimes that are in or close to <100 genes (**Fig. 3C,D** and **Table S9**). A small fraction of TE insertions with a higher frequency in selected populations were found in two of the studies. Taken together with the fact that there were very few differences in frequency of TE families, we propose that standing genetic variation presented by TEs plays a role in the evolution of aging, but it is unlikely to be a major driver of TE abundance differentiation. However, as we identified genomic locations of TEs only using PoPoolationTE2, which has been shown to have a low rate of false positives, we might miss insertions that would otherwise have been found by comparable software (Kofler et al. 2016; Nelson et al. 2017; Lerat et al. 2019).

Yet, we found that telomere maintenance, a key hallmark of aging known to be associated with mortality, diseases and the rate of senescence in several organisms might be improved in the late-breeding populations (Canela et al. 2007; López-Otín et al. 2013; Dantzer and Fletcher 2015; Foley et al. 2018; Whittemore et al. 2019). Among the three TE families constituting and maintaining *D. melanogaster* telomeres (Casacuberta 2017), *HeT-A* showed parallel increases in copy number in long-lived flies although the difference was less clear in two studies **(Fig. 3A** and **Fig. S1)**. Simultaneously, the few TEs transcriptionally upregulated in long-lived populations of Carnes2015 were almost exclusively telomeric elements (**Fig. 4B**). Despite similarities, the fundamental differences in telomeres between species make generalizations difficult (Mason et al. 2008). Moreover, previous studies in *D. melanogaster* and *C. elegans* failed to establish a connection between telomeres and lifespan, but telomere length might affect other traits such as fecundity (Raices et al. 2005; Walter et al. 2007). Also, in several species the rate of telomere shortening rather than the initial length itself was a better predictor for lifespan (Whittemore et al. 2019). Another complication yet to be addressed is if these patterns are caused by ‘intergenerational plasticity’ of telomere length, determined by paternal age at reproduction as observed in several mammals including humans (Eisenberg et al. 2012; Eisenberg and Kuzawa 2018). Thus, the exact impact of telomere length on evolutionary fitness and aging remains to be poorly understood.

### Is TE expression detrimental for longevity?

At the transcriptional level, age-dependent misregulation of TEs, thought to be resulting from a gradual decline in heterochromatin maintenance, has been proposed to be harmful for lifespan in *Drosophila* (Li et al. 2013; Chen et al. 2016; Wood et al. 2016; Brown and Bachtrog 2017; Guo et al. 2018), mice (De Cecco et al. 2019), and humans (Bogu et al. 2019). Further supporting the notion that expression of many TEs is detrimental, our RNA-seq analysis indicates that long-lived populations evolved to downregulate TE families, and this effect was even more apparent after we corrected for genomic copy numbers (**Fig. 4A,B** and **Table S18**). Considering the missing association between genomic abundance and TE transcription (**Fig. 4F**), this further suggests that lowering expression of TEs might be more important than purging them from the genome during the evolution of longevity.

Overall, however, TE expression appeared to be more strongly influenced by sex and age compared to selection regime. Interestingly, the trend of TEs being less expressed in late-breeding populations and upregulated with age was more pronounced in male flies, which further had generally higher levels of TE expression relative to females (**Fig. 4** and **Fig. S11A**). These findings are consistent with recent work showing that males suffer more from TE derepression during aging due to their entirely repetitive, heterochromatin-rich *Y* chromosome (Brown and Bachtrog 2017). However, if the divergent TE expression patterns between sexes are caused by differences in tissue compositions and whether this disparity explains sexual dimorphism in lifespan is yet to be confirmed. DNA-sequencing of male flies in the four experimental evolution studies would be necessary to determine if selection for postponed senescence had similarly strong effects on TE copy number of the *Y* chromosome.

### Did selection lead to differentiation in genes related to regulation of TE activity?

We also hypothesized that potential detrimental effects of TEs on longevity should be reflected by selection on genes related to TE regulation and transposition (**Fig. 5**). Although parallel genetic changes have been reported among the four studies (Fabian et al. 2018; Hoedjes et al. 2019), genetically and transcriptionally differentiated TE regulation genes were generally not shared between studies. Together with the missing functional enrichment associated with TE regulation, we hypothesize that improvement of chromatin structure/heterochromatin maintenance, piRNA-mediated silencing and modulators of transposition are not prime targets of selection during the evolution of longevity. This, however, does not preclude that other means of TE protection have evolved. It is becoming increasingly evident that TE expression acts as a causative agent of inflammation and immune activation in mammals (Kassiotis and Stoye 2016; De Cecco et al. 2019). Interestingly, Carnes2015, Fabian2018, and Remolina2012 all found significant divergence in innate immunity genes, whereas Fabian et al. (2018) demonstrated an improved survival upon infection and alleviated immunosenescence in the long-lived populations. Rather than reducing TE copy number and expression, selection might preferentially act on immunity genes to reduce TE-mediated inflammation and increase tolerance to TEs with extended lifespan as a consequence. It remains to be explored to what degree innate immune pathways other than the RNAi machinery contribute to TE regulation in *D. melanogaster*.

### Is reproduction at old age associated with an increased TE content?

Our findings suggest that neither genetic drift nor pervasive selection on TEs or genes related to TE regulation are predominant drivers of the differences in TE abundance. The most parsimonious explanation for our results therefore is that postponed reproduction increases the chance for many TEs to be inserted into the germline and passed on to the next generation. In particular, TE families of high frequency which are putatively low in transpositional activity might need the prolonged chronological time offered by late-life reproduction to achieve a successful genomic insertion (**Fig. 2**). Over many generations, flies breeding at old age would have accumulated more TEs in the genome than populations reproducing early in life. Supporting this hypothesis, it has been demonstrated that most TE families had a higher rate of insertions in the ovaries of older relative to young *P-element* induced dysgenic hybrids, even though at the same time fertility was restored and improved with age (Khurana et al. 2011). However, if this applies to non-dysgenic fruit flies and whether it can result in a larger number of TEs over multiple generations has to our knowledge not yet been observed. Thus, TE accumulation in late-breeding populations is comparable to the regularly observed positive correlation between parental age and number of *de novo* mutations in offspring (Goldmann et al. 2019; Sasani et al. 2019). In line with this, genome-wide measures of nucleotide diversity were also repeatably larger in late-breeding populations across four experiments (**Table S7**). Although, we have not ruled out that greater nucleotide diversity was driven by genetic drift or balancing selection as proposed by one study (Michalak et al. 2017).

Opposing our hypothesis, two recent studies in termites (Elsner et al. 2018) and *D. melanogaster* (Erwin and Blumenstiel 2019) suggest that the germline is protected from TE invasions through increased transcription of the piRNA machinery. Indeed, our expression analysis confirms that many genes associated with transcriptional and post-transcriptional TE silencing tend to be upregulated with age. Despite this, many TE families had a higher copy number in populations reproducing late in life. It therefore remains to be determined whether this age-dependent upregulation of TE regulation genes really equates to reduced insertional activity, since potential and realized TE repression might not necessarily match. The observation that these genes also tended to be more expressed in controls relative to selected flies in Carnes2015 further poses the question whether there is a trade-off between TE silencing in the germline and lifespan, which could be another mechanism explaining the rising TE abundance in the genomes of long-lived flies.

Altogether, our work presents a novel viewpoint on the poorly understood role of TEs in aging and longevity that is largely, but not exclusively, neutral. However, the caveat remains that we are unable to rule out that survival of selected populations would be further extended if they had a reduced TE content and expression. In-depth studies tracking piRNA production in the germline together with direct measures of TE transposition rates throughout life or measuring longevity upon knockdown and overexpression of TEs would be crucial experiments to obtain a more complete picture.

## MATERIALS AND METHODS

### Datasets

We utilized genomic data from four independent studies performing laboratory selection for postponed reproduction on wild-derived replicate populations by only allowing flies of relatively old age to contribute to subsequent generations, whereas controls reproduced early in life (Remolina et al. 2012; Carnes et al. 2015; Fabian et al. 2018; Hoedjes et al. 2019) (**Table S1**). The experimental designs of the studies were overall comparable, but notable differences include the mode of selection, maintenance of controls, variable source populations, number of replicate populations and generations at the time of sequencing. Moreover, Hoedjes2019 performed the selection for postponed senescence on three varying larval diets ranging from low to high sugar/protein content. The genomic analysis was based on available raw fastq files from whole-genome pool-sequencing of 100 to 250 females. RNA-seq data from Carnes et al. (2015) consisted of raw fastq files from pools of 50 flies. The study included transcriptomes of all selected and control populations, for which both sexes at two ages 3-5 days (young) and 26-35 days of age (old) have been sequenced in replicates. Microarray expression data from Remolina et al. (2012) are derived from heads and abdomens from females at the age of 1, 5, 15, 30, and 50 days of age from the three control and selected populations. See methods in the publications of each study for details on experimental design and **Table S1** for a summary. For simplicity, we refer to Carnes et al. (2015) as Carnes2015, Fabian et al. (2018) as Fabian2018, Hoedjes et al. (2019) as Hoedjes2019, and Remolina et al. (2012) as Remolina2012 throughout this report. All statistics were done in *R* using in-built functions unless otherwise stated. More details on the bioinformatic pipeline are available in **Table S20**.

### Genome-wide TE abundance

To quantify the number of genomic insertions for each TE family in selected and control populations we used DeviaTE (Weilguny and Kofler 2019) (**Table S20**). In brief, DeviaTE maps raw reads to an incorporated library of 179 TE family consensus sequences (Sackton et al. 2009; Bergman et al. 2018) and normalizes the obtained coverage values by the average depth of the same five single-copy genes (**Fig. S2**). The distribution of normalized values reflects fluctuations in insertion number estimates within a TE family, where averaging over all consensus positions of a TE family gives the mean abundance per haploid genome (see Weilguny and Kofler 2019 for details). We restricted our downstream analysis to TE families that had a study-average of >=0.5 insertions for at least 80% of the consensus positions within a TE family sequence (**Fig. S3**). Thus, we excluded all families with very low abundance and potentially wrongly mapped reads, and TE families of which only small fractions of the whole consensus sequence were covered.

We then investigated if TE families vary in genomic abundance between control and selected populations using three different approaches (see **Fig. S1** for a comprehensive description). In our least conservative approach #1, we analyzed studies by fitting *Regime* (control, selected) and *Population[Regime]* (replicate populations nested within regime) to normalized coverage values of consensus sequence positions within a TE family. For Hoedjes2019, we used a different model and included *Regime, Diet* (low, medium, high protein/sugar larval diet regime), and the *Regime x Diet* interaction. To correct for multiple testing, we applied a Bonferroni cut-off at *α* = 0.01. We further used *SuperExactTest* (Wang et al. 2015) to analyze if the overlap of TEs with a significantly higher genomic abundance in selected (“S>C”) or control populations (“C>S”) between postponed senescence studies is expected by chance. The normalized coverage values were averaged to obtain a single insertion estimate per TE family and population, and these values used for all the remaining analyses.

For approach #2, we arcsine square root transformed proportions of TE family copy number relative to the total genomic TE content within a population and analyzed all studies together rather than independently by fitting *Study* (four levels: Carnes2015, Fabian2018, Hoedjes2019, Remolina2012), *Regime* and the *Study x Regime* interaction as factors. TE families with an FDR < 0.05 were considered significant.

Finally, our approach #3 is the most conservative as we only considered TE families that showed a consistent increase or decrease in copy number (i.e. average of insertion estimates across all consensus positions) within all selected relative to all control populations in each study and within diet regimes of Hoedjes2019.

To analyze differences in the total genomic and subclass-specific (LTR, non-LTR, TIR) TE content, we summed up all TE insertion estimates within a population and fit models with *Study, Regime* and the *Study x Regime* interaction.

### Genomic TE locations and activity/age of TE families

We first masked the *D. melanogaster* reference (v.6.27) for TEs present in the DeviaTE library using RepeatMasker (Smit et al. 1996) (**Table S20**). We then trimmed reads with cutadapt (Martin 2011) and mapped them using bwa bwasw (Li and Durbin 2009). PoPoolationTE2 was then employed to obtain the exact genomic positions and population frequency of TE insertions on chromosomes *X, 2, 3*, and *4* of each study using the joint analysis mode, which finds insertions by combining all samples rather to considering them separately (Kofler et al. 2016). Importantly, while TE abundance is quantified by the total number of reads mapping to a TE relative to single-copy genes (Weilguny and Kofler 2019), identifying the exact genomic location of insertions requires mates of a read-pair to map discordantly to the reference genome and TE sequence, and strongly depends on the sequencing depth and number of populations (Cridland et al. 2013; Kofler et al. 2016; Lerat et al. 2019). For each TE family, we calculated the average population frequency across all of its detected genomic locations within a population as a proxy for active or recent transposition events and evolutionary age (Kofler et al. 2015). We used Spearman’s correlation analysis to compare average frequency values of each study to average frequencies from a natural South African (SA) population sequenced to a high genomic coverage (Kofler et al. 2015), and to correlate TE abundance with average frequency. We employed t-tests to analyze if average population frequency from the SA population varies between TE families more abundant in selected or control populations, and also performed this analysis using only the top10 TEs with the largest log_2_ FC values of abundance change.

### Genome-wide nucleotide diversity and genetic drift simulations

We mapped trimmed paired-end reads against the repeat-masked reference genome, the TE library from DeviaTE (Weilguny and Kofler 2019), *Wolbachia pipientis* (NC_002978.6), and two common gut bacteria *Acetobacter pasteurianus* (AP011121.1), and *Lactobacillus brevis* (CP000416.1) using bwa *mem* (Li and Durbin 2009), and removed duplicates using PicardTools (**Table S20**). We then filtered and created pileup files using samtools *mpileup* (Li et al. 2009). To calculate nucleotide diversity *π* and Watterson’s *θ* across non-overlapping 100kb windows we used Popoolation (Kofler et al. 2011) and then fitted ANOVA models including the factors *Chromosome* (*X, 2L, 2R, 3L, 3R, 4*), *Diet, Regime*, and the *Diet x Regime* interaction for Hoedjes2019, and *Population[Regime], Chromosome*, and *Regime* for all other studies. Average coverage across major chromosomal arms was 162x, 101x, 41x, and 23x for Fabian2018, Hoedjes2019, Remolina2012, and Carnes2015, respectively. We detected reads mapping to the genome of the intracellular bacterium *Wolbachia* in all populations. To test if TE family abundance differences can be caused by genetic drift alone, we compared proportions of S>C and C>S TEs from 5,000 simulations of TE frequency change to observed proportions from approach #1 and #3. See Supplementary Methods for more details.

### TE frequency differences

To identify genomic TE insertion sites putatively involved in lifespan and aging, we analyzed differences in arcsine square root transformed insertion frequencies between selected and control populations fitting models with *Regime* for Carnes2015, Fabian2018 and Remolina2012, and with factors *Diet, Regime*, and *Diet x Regime* for Hoedjes2019. Bonferroni correction at *α* = 0.05 was used to correct for multiple testing. Functional annotations were supplemented using SnpEff (v.4.0e, Cingolani et al. 2012) considering TE insertions within 1000 bp of the 5’ and 3’ UTR as upstream or downstream of a gene.

We further analyzed if each TE family varies in frequency between regimes by fitting the factors of *Diet, Regime, and Diet x Regime* for Hoedjes2019, or *Regime* and *Population[Regime]* for all other studies on arcsine square root transformed insertion site frequencies. FDR values were obtained by using “p.adjust” in *R* and TE families considered significant at FDR < 0.05.

### RNA-seq analysis

RNA-seq data from Carnes et al. (2015) consisted of two replicates of young and old males and females from all control and selected populations (**Table S1**). Raw reads were filtered using cutadapt (Martin 2011) and mapped to the repeat-masked reference genome, the TE library from DeviaTE, *Wolbachia pipientis, Acetobacter pasteurianus*, and *Lactobacillus brevis* (see above) using STAR (Dobin et al. 2013) (**Table S20**). Read counts were obtained using featureCounts (Liao et al. 2013). We next pre-filtered read count data by excluding all genes and TEs that did not have a sum of 400 counts across all 80 samples (i.e. on average 5 counts per sample). Five TE families that are not known to occur in *D. melanogaster* passed this filter and were excluded. For simplicity, the analysis was performed on average read counts from two replicates, as all replicates were highly significantly correlated (Pearson’s r ranging from 0.95 to 1, significant after Bonferroni correction). To analyze differential expression, we fit models using read counts of genes and TEs with DESEq2 in *R* (Love et al. 2014). First, a model testing the main effects of *Regime* (selected vs control), *Sex* (male vs female), and *Age* (young vs old) was fit. As the sex term was significant for most TEs, we decided to analyze males and females separately and fitted models with *Regime* and *Age* to analyze the main effects. To examine the interaction, we also fitted models including *Regime x Age*. We obtained log_2_ fold change values for each factor and the library-size normalized read counts from DEseq2 for further analysis. To investigate average expression per TE insertion, we divided read counts of TEs from females by the number of genomic insertions observed in each population, assuming that genes and 13 TEs that did not pass our filters in the genomic analysis have a single copy in the genome.

### Evolution of TE regulation genes

The list of genes involved in TE regulation consisted of piRNA pathway genes also analyzed in Erwin and Blumenstiel 2019 and Elsner et al. 2018, and genes involved in heterochromatic and chromatin structure from Lee and Karpen 2017. We further added 7 genes involved in these functions, and genes annotated to “regulation of transposition” (GO:0010528) and “transposition” (GO:0032196) according to FlyBase so that we ended up with a total of 96 genes (**Table S19**). We then screened the published genomic candidate gene lists from Carnes2015, Fabian2018, Hoedjes2019 and Remolina2012 for these genes. We also compared TE regulation genes to differentially expressed genes from the RNA-seq analysis of Carnes2015 (see above). We further obtained normalized microarray expression data from Remolina2012 of female flies at 1, 5, 15, 30, and 50 days of age (**Table S1**). Notably, the expression data were created from flies at 40 generations of selection compared to 50 generations in the genomic analysis. We fit a mixed effects model similar to the one used in their original publication with *Age, Regime*, and *Age x Regime* as fixed and *replication within population-age combination* as random effect. The two available tissues (heads and abdomens) were analyzed separately. A gene was considered to be differentially expressed if it had an FDR < 0.05 unless otherwise stated.

## DATA ACCESSIBILITY

Accession numbers to the raw genomic and transcriptomic data can be found in **Table S1** and in the original studies (Remolina et al. 2012; Carnes et al. 2015; Fabian et al. 2018; Hoedjes et al. 2019). RNA-seq data were obtained directly from the authors (Wen Huang and Trudy Mackay, active download links in supplementary code on GitHub). Scripts to all analyses and raw output files are available at: https://github.com/FabianDK/LongeviTE. Additional raw output and edited files used to analyze TE abundance and nucleotide diversity, results for the microarray analysis, and boxplots showing the number of insertions for significant TE families are available on Dryad (DOI: 10.5061/dryad.s7h44j13r).

## ACKNOWLEDGEMENTS

We are grateful to Alexis Braun, three anonymous reviewers and the associate editors for comments on the manuscript. We also thank Robert Kofler, Andrea Betancourt, Frank Jiggins and Lukas Weilguny for helpful discussions. This work was supported by the Wellcome Trust (WT098565/Z/12/Z to J.M.T. and L.P.), EMBL (H.M.D. and J.M.T.), and Comisión Nacional de Investigación Científica y Tecnológica – Government of Chile (CONICYT scholarship to M.F.). We are further grateful for financial support from the Society for Molecular Biology & Evolution enabling us to present this work at the annual meeting (SMBE 2019, Carer travel award & registration award to D.F.).

