## Supplementary Material for "Transposable element landscape in *Drosophila* populations selected for longevity"

SUPPLEMENTARY RESULTS

#### TE abundance differences between regimes using three approaches

We used three different approaches to determine whether TE families vary in copy number between control and selected populations (**Fig. S1**, **Table S2**). In our least stringent approach (approach #1), we analyzed the average differences between regimes in each study and performed statistics using the complete set of insertion estimates at each position in the TE family consensus sequence (see main Results text).

In our second approach (approach #2), we first averaged normalized coverage values for each TE family to obtain insertion estimates. Because the total number of TEs was significantly different between studies (ANOVA $P < 0.0001$ of Study factor; **Fig. 1C** and **Table S6**), we normalized the insertion counts of each individual TE family by the total insertions in the corresponding population and analyzed all four studies combined using arcsine square root transformed proportions (**Fig. S1B** and **Fig. S1C**). At an FDR of $<0.05$, we identified 41 TEs with a significant regime factor of which 34 were more and 7 less abundant in selected populations, confirming our findings of individually analyzed studies using approach #1 (**Table 1**, **Table S2**). Comparable tendencies were obtained when we analyzed the raw insertion counts or standardized z-scores of insertions and changing the significance threshold did not alter the fact that S>C TEs were more frequent than C>S (not shown).

Finally, our most conservative analysis (approach #3), identified TEs with consistent differences in genomic TE insertions between all control and all selected populations (**Table 1, Table S2**). The number of TEs with abundance differences were generally smaller compared to approach #1 indicating that TEs within control or selected populations do not always behave in a parallel way. The insertion tendency in favor of S>C was again apparent in three studies but varied between larval diet conditions in Hoedjes2019: selection for postponed reproduction resulted in a clear tendency of more S>C than C>S TEs when flies were selected on a low sugar/protein diet. This trend was less pronounced for populations selected on the medium diet, while we observed the opposite pattern for the high sugar/protein condition (i.e. more C>S than S>C TEs). Qualitatively similar results were obtained when we used approach #1 to analyze differences between long-lived and control populations in

Hoedjes2019 separately for each diet (**Fig. S4, Table S2** and **S3**). Yet, the overall increase in TE insertions in late-breeding populations of Hoedjes2019 was apparent in our model which corrects for the effect of diet (**Fig. 1** and **Table 1**).

To analyze and visualize differences in TE abundance, we obtained δinsertion values by subtracting average insertions across controls from copy numbers in selected populations for each TE family (as in upper panel of **Fig. S4** and **Fig. S5A**). As expected, we found that δinsertion values scaled with genomic copy number of TE families (Pearson's r, Carnes2015: 0.79, Fabian2018: 0.51, Hoedjes2019: 0.77, Remolina2012: 0.6, all $P < 0.0001$), we also obtained $\log_2$ fold change (FC) values by dividing average insertions of selected by control populations for each TE (**Fig. 1A** and lower panel of **Fig. S4**).

### SUPPLEMENTARY METHODS

#### Genetic drift simulations

As the analyzed studies did not include data from the ancestral populations, we used average TE frequencies from the South African population (Kofler et al. 2015) as a starting point in our simulations. To simulate frequency change of each TE, we set population sizes (N), generation times and number of replicate populations as mentioned in the original publications (**Table S1**). As some studies reported a range of population sizes, we performed simulations using the lower and upper limits. We used the *rbinom* function in *R* and drew from a size of 2N considering the diploid genome and given the average frequency of a TE as probability to be drawn. The number of successful draws was divided by 2N to obtain the new TE frequency, which was then used as input for the next draw. We repeated this process until the generation times at sequencing of control and selected populations were reached. Using the simulated TE frequencies in the last generation, we calculated average TE frequency across all simulated replicates within breeding regime. Next, we obtained the proportion of TEs with a higher (S>C) and lower frequency (C>S) in selected populations and calculated the $\log_2$ relative proportion (i.e. $\log_2$ of S>C proportion divided by C>S proportion). As an additional approach, we also acquired the $\log_2$ relative proportion by using the number of TEs with consistent differences in frequency between all the populations of the regimes instead of averaging. We performed 5,000 simulations to get a distribution of relative proportions expected under genetic drift. P-values were calculated by dividing the number of simulations that resulted in a larger or equal proportion as observed in the actual analysis by the total number of simulations.

#### SUPPLEMENTARY TABLES

**Table S1.** Experimental designs of evolution for postponed senescence studies. N, number of populations per selection regime. Age at egg collection, corresponds to the age in days at which eggs were collected to establish the next generation. Lifespan from mated individuals as reported in the corresponding publications, except for Hoedjes et al. 2019, for which lifespan was averaged by manually extracting data from Figure 4 in May et al. 2019 (data from different generation as whole genome sequences). SE, single-end reads. PE, paired-end reads. SRA, Sequence Read Archive accession number

| Study | Regime | N | Age at egg collection | Population size | Generations | Lifespan in days | Pool-size & meta-data | Seq-type | SRA |
|---|---|---|---|---|---|---|---|---|---|
| Carnes et al. (2015) | Selected | 5 | Progressive increase starting from 28 days, after generation 15: 70 – 84 days | 1000 - 2000 | 170 | Mean of males & females: 62.1 | **DNA:** 100 females **RNA:** 50 whole males/females at 3-5 (young) and 26-35 days (old), 2 replicates | **DNA:** 68-72 bp, PE **RNA:** 125 bp, SE | **DNA:** PRJNA286855 **RNA:** Obtained from Wen Huang / Trudy Mackay |
| | Controls | 5 | 14 days | 1000 - 2000 | 850 | Mean of males & females: 35.7 | | | |
| Fabian et al. (2018) | Selected | 4 | Progressive increase starting from 22 days; after generation 25: 20 - 30% longest surviving | 300 | 144 - 148 | Median: Males: 72 Females: 62.5 | 100 females | 100 bp, PE | PRJEB28048 |
| | Controls | 2 | Randomly chosen between 4 and 30 days | 300 | 293, 310 | Median: Males: 48 Females: 45 | | | |
| Hoedjes et al. (2019) | Selected | 12 | 16 – 18 days | 2000 - 4000 | 58 | Mean: Males: ~62 Females: ~61 | 250 females | 100 bp, PE | PRJNA564570 |
| | Controls | 12 | 2 – 4 days | 2000 - 4000 | 115 | Mean: Males: ~57 Females: ~51 | | | |
| Remolina et al. (2012) | Selected | 3 | Progressive increase starting from 28 days, up to 40 days | 110-160 mating pairs (i.e. 50% less than controls; 220-320 individuals) | **DNA:** 50 **RNA:** 40 | Mean: Males: 44.2 Females: 34.1 | **DNA:** 100 females **RNA:** 7 females, heads & abdomens, 5 ages from 1 to 50 days, 2 replicates | **DNA:** 74 bp, PE 6 lanes/sample **RNA:** Microarray | **DNA:** PRJNA185744 **RNA:** GSE38106 |
| | Controls | 3 | 14 days | 220-320 mating pairs (i.e. 440-640 individuals) | ~80 | Mean: Males: 40.7 Females: 29.9 | | | |

**Table S2 (Excel File).** Summary statistics for TE abundance differences using three different approaches. Sheets labelled 'Approach_#1&3' show genome-wide mean TE family abundance estimates from DeviaTE (Weilguny and Kofler 2019) for each population and regime within the four studies. Populations are labelled according to selection regime and their published names: Cont, early-breeding controls; Sel, late-breeding selection regime. For Hoedjes2019: M, medium sugar/protein; H; high sugar/protein; L, low sugar/protein larval diet; E, early reproduction; P, postponed reproduction. Columns 'Diff_SelCont' and 'log2_SelCont' denote the difference in average abundance between selection and control regimes, and the log2 fold change. Summary of statistical models is shown in columns denoted as 'Df' (degrees of freedom); 'F_value'; 'P_value', 'Bonf' (TRUE if TE family passed Bonferroni corrected P-value cut-off for regime factor), followed by the corresponding factor in the fitted models ('Regime' is selection vs control; 'nested' is population nested within regime; 'Diet' is low, control, high sugar/protein diet conditions). The columns 'ConsSC' and 'ConsCS' correspond to approach #3 and indicate whether a TE family was consistently more abundant in all selected populations relative to controls (SC), or more abundant in all control relative to selected populations (SC) within a study. The sheet 'Approach_#2' includes summary statistics for the model that analyzed TE family abundance normalized by total genomic content combining all four studies. For more details, see **Fig. S1** and Materials and Methods.

**Table S3.** Number of detected and significant TE families per larval diet of Hoedjes2019 with percentage of TEs more abundant in selected (S>C) or controls (C>S) or not different between regimes (n.s.) from models analyzing diets separately (approach #1, **Fig. S1**). Summary of statistics can be found in **Table S2**.

| Diet | N | N (sign.)[a] | S>C | C>S | n.s. |
|---|---|---|---|---|---|
| Low | 115 | 108 | 90% | 5% | 4% |
| Medium | 115 | 99 | 61% | 27% | 12% |
| High | 115 | 98 | 6% | 81% | 13% |

[a]Significant after Bonferroni correction at $\alpha$=0.01.

**Table S4 (Excel File). Summary statistics for TE abundance differences considering only the last 200 bp at the 3'-ends of TE family consensus sequences.** Sheet 'Summary' gives an overview of significant TE families across the three approaches. Sheets labelled 'Approach_#1&3' show genome-wide mean TE family abundance estimates from DeviaTE for each population and regime within the four studies. Populations are labelled according to selection regime and their published names: Cont, early-breeding controls; Sel, late-breeding selection regime. For Hoedjes2019: M, medium sugar/protein; H; high sugar/protein; L, low sugar/protein larval diet; E, early reproduction; P, postponed reproduction. Columns 'Diff_SelCont' and 'log2_SelCont' denote the difference in average abundance between selection and control regimes, and the log2 fold change. Summary of statistical models is shown in columns denoted as 'Df' (degrees of freedom); 'F_value'; 'P_value', 'Bonf' (TRUE if TE family passed Bonferroni corrected P-value cut-off for regime factor), followed by the corresponding factor in the fitted models ('Regime' is selection vs control; 'nested' is population nested within regime; 'Diet' is low, control, high sugar/protein diet conditions). The columns 'ConsSC' and 'ConsCS' correspond to approach #3 and indicate whether a TE family was consistently more abundant in all selected populations relative to controls (SC), or more abundant in all control relative to selected populations (SC) within a study. The sheet 'Approach_#2' includes summary statistics for the model that analyzed TE family abundance normalized by total genomic content combining all four studies. For more details, see **Fig. S1** and Materials and Methods.

**Table S5 (Excel File). Summary statistics for sequence abundance differences.** We summed up normalized coverage values across all consensus sequence positions within a TE family to obtain sequence abundance and then analyzed differences between control and selection regimes using approach #2 and #3. Sheet 'Summary' gives an overview of significant TE families across the three approaches. Populations are labelled according to selection regime and their published names: Cont, early-breeding controls; Sel, late-breeding selection regime. For Hoedjes2019: M, medium sugar/protein; H; high sugar/protein; L, low sugar/protein larval diet; E, early reproduction; P, postponed reproduction. Columns 'Diff_SelCont' and 'log2_SelCont' denote the difference in average abundance between selection and control regimes, and the log2 fold change. Summary of statistical models is shown in columns denoted as 'Df' (degrees of freedom); 'F_value'; 'P_value', 'FDR' (false discovery rate), followed by the corresponding factor in the fitted model. The columns 'ConsSC' and 'ConsCS' correspond to approach #3 and indicate whether a TE family was consistently more abundant in all selected populations relative to controls (SC), or more abundant in all control relative to selected populations (SC) within a study. For more details, see **Fig. S1** and Materials and Methods.

**Table S6 (Excel File). Differences in total TE content.** Total TE content per population was obtained by summing up all copy numbers from each TE family (see **Table S2**). Sheet 'SummaryStat' shows the summary of ANOVA model analyzing variation in total insertions for the effects of study, regime and the interaction factor. Cont, early-breeding controls; Sel, late-breeding selection regime. Columns 'Diff_SelCont' and 'log2_SelCont' denote the difference in average total content between selection and control regimes, and the $\log_2$ fold change. Summary of statistical models is shown in columns denoted as 'Df' (degrees of freedom); 'F_value'; 'P_value' for each factor. Sheet 'Total_TE_Insertions' shows the total TE insertions for each population. Also see **Fig. 1D** and **Fig. S6**.

**Table S7.** Differences in nucleotide diversity estimators between breeding regimes (Control and Selected) given as genome averages and standard error of the mean. *F* and P-values correspond to factors used in models, which differ between Hoedjes2019 and the other studies.

| Study | Estimator | Control Mean | Selected Mean | *F* Regime | P Regime | *F* Chrom.[a] | P Chrom.[a] | *F* Pop[Regime][a] | P Pop[Regime][a] | *F* Diet | P Diet | *F* Regime x Diet | P Regime x Diet |
|---|---|---|---|---|---|---|---|---|---|---|---|---|---|
| Carnes2015 | π | 0.00242 (± 2.3e-05) | 0.00332 (± 2.3-05) | $F_{1,11284}$=976.8 | <2.2e-16 | $F_{5,11284}$=570.1 | <2.2e-16 | $F_{8,11284}$=18.3 | <2.2e-16 | - | - | - | - |
| | θ | 0.00230 (± 2.1e-05) | 0.00294 (± 1.9e-05) | $F_{1,11284}$=643.5 | <2.2e-16 | $F_{5,11284}$=594.2 | <2.2e-16 | $F_{8,11284}$=31.3 | <2.2e-16 | - | - | - | - |
| Fabian2018 | π | 0.00148 (± 3.2e-05) | 0.00152 (± 2.5e-05) | $F_{1,6863}$=1.8 | 0.179 | $F_{5,6863}$=304 | <2.2e-16 | $F_{4,6863}$=616.6 | <2.2e-16 | - | - | - | - |
| | θ | 0.00113 (± 1.8e-05) | 0.00132 (± 1.6e-05) | $F_{1,6863}$=89.8 | <2.2e-16 | $F_{5,6863}$=314.7 | <2.2e-16 | $F_{4,6863}$=1196.2 | <2.2e-16 | - | - | - | - |
| Hoedjes2019 | π | 0.00393 (± 1.5e-05) | 0.00408 (± 1.5e-05) | $F_{1,27535}$=61.9 | <2.2e-16 | $F_{5,27535}$=1505.2 | <2.2e-16 | - | - | $F_{2,27535}$=12.7 | <3.2e-6 | $F_{2,27535}$=21 | <7.7e-10 |
| | θ | 0.00333 (± 1.2e-05) | 0.00346 (± 1.2e-05) | $F_{1,27535}$=74.3 | <2.2e-16 | $F_{5,27535}$=1722.9 | <2.2e-16 | - | - | $F_{227535}$=38 | <2.2e-16 | $F_{2,27535}$=28 | <7.5e-13 |
| Remolina2012 | π | 0.00335 (± 3e-05) | 0.00394 (± 3.2e-05) | $F_{1,6866}$=234.2 | <2.2e-16 | $F_{5,6866}$=400.2 | <2.2e-16 | $F_{4,6866}$=3.8 | 0.004 | - | - | - | - |
| | θ | 0.00283 (± 2.4e-05) | 0.00346 (± 2.7e-05) | $F_{1,6866}$=396.1 | <2.2e-16 | $F_{5,6866}$=397.2 | <2.2e-16 | $F_{4,6866}$=2.2 | 0.062 | - | - | - | - |

[a]Chrom., Chromosome. Pop[Regime], population nested within regime.

**Table S8 (Excel Sheet)**. **Shared TE families across four independent studies.** We analyzed study-overlaps of TE families with a higher (S>C) or lower genomic abundance (C>S) in the selected populations. To test if overlaps are expected by chance, we used *SuperExactTest* in R.

**Table S9 (Excel File). Frequency differentiation of individual TE insertions.** We identified the exact genomic positions and frequency of TEs in Carnes2015, Fabian2018, Hoedjes2019 and Remolina2012 using PoPoolationTE2. To identify candidate insertions, we fitted models with the Regime term (for Hoedjes2019 only: included Diet and Regime x Diet interaction) to arcsine square root transformed insertion frequencies and adjusted P-values using Bonferroni correction. We did not identify any significant TE insertions in Carnes2015 and Remolina2012, but in Fabian2018 and Hoedjes2019 (shown in sheets labelled 'Candidates'). TE frequency of control and selected populations are labelled 'Cont' and 'Sel'. In Hoedjes2019, populations are named LE, LP, ME, MP, HE, and HP, where 'L/M/H' denotes if selection took place on larval diet with low/medium/high sugar and protein content; while 'E/P' indicates whether the population was early/late reproducing. We also provide average frequencies across all populations, controls, and selected populations ('AvFreq', 'AvFreqC', 'AvFreqS'). Annotations and effects of TE insertions were supplemented using SnpEff. If a TE insertion was 1kb of the gene boundaries, it was considered to be upstream or downstream. Statistics for all tested insertions are in the sheets labelled 'All insertions'.

**Table S10.** Average number of insertions (Ins.), average frequency (Freq.), and differences between selection and control regimes (δIns. and δFreq.) for TEs with a significant difference in abundance and frequency. Summary statistics are available in **Table S2** for copy number and **Table S11** for TE family frequency differences.

| | TE | Ins. (S) | Ins. (C) | Freq. (S) | Freq. (C) | δIns. | δFreq. | Same Direction[a] |
|---|---|---|---|---|---|---|---|---|
| **Carnes2015** | *1360* | 96.80 | 78.07 | 0.75 | 0.64 | 18.73 | 0.11 | yes |
| | *baggins* | 27.73 | 16.82 | 0.70 | 0.51 | 10.91 | 0.19 | yes |
| | *Circe* | 35.39 | 24.44 | 0.71 | 0.49 | 10.95 | 0.21 | yes |
| | *diver2* | 7.37 | 5.67 | 0.85 | 0.67 | 1.70 | 0.18 | yes |
| | *S-element* | 21.61 | 15.16 | 0.74 | 0.60 | 6.45 | 0.14 | yes |
| | *Cr1a* | 79.90 | 64.72 | 0.66 | 0.56 | 15.18 | 0.10 | yes |
| | *GATE* | 13.02 | 10.48 | 0.69 | 0.49 | 2.54 | 0.19 | yes |
| | *Rt1a* | 9.14 | 6.88 | 0.94 | 0.73 | 2.26 | 0.22 | yes |
| | *Max-element* | 12.01 | 11.43 | 0.79 | 0.53 | 0.58 | 0.26 | yes |
| | *I-element* | 35.12 | 66.81 | 0.80 | 0.58 | -31.69 | 0.22 | no |
| | *297* | 51.27 | 54.15 | 0.70 | 0.45 | -2.88 | 0.25 | no |
| | *Doc2-element* | 21.04 | 28.64 | 0.89 | 0.60 | -7.61 | 0.29 | no |
| | *Doc3-element* | 26.52 | 27.70 | 0.81 | 0.59 | -1.18 | 0.22 | no |
| | *Doc4-element* | 4.65 | 3.31 | 0.81 | 0.45 | 1.34 | 0.35 | yes |
| | *Porto1* | 14.46 | 18.73 | 0.68 | 0.45 | -4.27 | 0.22 | no |
| | *Fw2* | 4.05 | 3.20 | 0.92 | 0.59 | 0.85 | 0.33 | yes |
| | *G3* | 2.40 | 1.93 | 0.72 | 0.43 | 0.48 | 0.29 | yes |
| | *G4* | 13.32 | 10.08 | 0.88 | 0.64 | 3.24 | 0.24 | yes |
| | *G5* | 6.83 | 5.59 | 0.83 | 0.62 | 1.25 | 0.21 | yes |
| | *gypsy10* | 5.22 | 3.43 | 0.80 | 0.53 | 1.80 | 0.27 | yes |
| | *gypsy7* | 7.45 | 5.58 | 0.86 | 0.46 | 1.88 | 0.41 | yes |
| | *gypsy9* | 1.70 | 1.46 | 0.85 | 0.62 | 0.24 | 0.23 | yes |
| | *invader1* | 13.32 | 10.73 | 0.80 | 0.45 | 2.59 | 0.35 | yes |
| | *invader2* | 30.92 | 32.49 | 0.75 | 0.59 | -1.57 | 0.16 | no |
| | *mariner2* | 9.67 | 6.62 | 0.74 | 0.56 | 3.05 | 0.18 | yes |
| | *accord2* | 7.79 | 8.96 | 0.93 | 0.70 | -1.17 | 0.23 | no |
| | *Quasimodo* | 45.96 | 34.90 | 0.74 | 0.49 | 11.05 | 0.25 | yes |
| | *rooA* | 11.66 | 6.43 | 0.75 | 0.49 | 5.23 | 0.26 | yes |
| | *X-element* | 13.35 | 9.62 | 0.50 | 0.25 | 3.73 | 0.25 | yes |
| | *Rt1b* | 24.91 | 21.62 | 0.70 | 0.45 | 3.29 | 0.25 | yes |
| | *S2* | 5.94 | 3.15 | 0.78 | 0.46 | 2.79 | 0.32 | yes |
| | *Tc1* | 21.42 | 14.10 | 0.66 | 0.37 | 7.32 | 0.29 | yes |
| | *Tc1-2* | 18.33 | 11.97 | 0.78 | 0.60 | 6.36 | 0.18 | yes |
| | *Tc3* | 4.32 | 3.06 | 0.91 | 0.59 | 1.26 | 0.32 | yes |
| **Fabian2018** | *P-element* | 9.95 | 19.02 | 0.03 | 0.07 | -9.07 | -0.04 | yes |

| | | | | | | | | |
|---|---|---|---|---|---|---|---|---|
| **Hoedjes2019** | *roo* | 69.61 | 64.53 | 0.04 | 0.04 | 5.08 | 0.00 | yes |
| | *jockey* | 23.52 | 23.24 | 0.03 | 0.03 | 0.28 | 0.00 | yes |
| | *P-element* | 20.50 | 20.00 | 0.03 | 0.03 | 0.49 | 0.00 | yes |
| **Remolina2012** | No TE type significant for frequency differences. | | | | | | | |

[a]Same Direction denotes TEs that show the same sign for $\delta$Ins. and $\delta$Freq.

**Table S11 (Excel File). Frequency differentiation of TE families.** We tested if TE families show frequency differentiation between control (Cont) and selected populations (Sel) by fitting the effects of Regime and Population[Regime] for Carnes2015, Fabian2018, and Remolina2012, or Regime, Diet, and the Regime x Diet interaction for Hoedjes2019 to arcsine square root transformed insertion frequencies. P-values were adjusted using FDR. Columns dFreq and log2Freq show the difference and log2 fold change between selected and control regimes. NAs indicate TE families with only a single detected insertion for which models could not be fit. TE insertion frequencies used for this analysis can be found in **Table S9**.

**Table S12.** Number of differentially expressed TE families and genes in parentheses at an adjusted P-value of <0.05. We fit different models to analyze the effects of Regime (C, control; S, selected), Age (young; old), and Sex (M, males; F, females). ‘RxA’ refers to the Regime x Age interaction. Sample sizes varied between models and factors dependent on auto-filtering by DESeq2, and ranged from 107 to 123 TEs and 9,896 to 13,249 genes. The median sample size across all models and factors was n = 122 for TEs and n = 13,078 for genes. DESeq2 outputs for all models can be found in **Table S13**.

| Model | Regime | Age | Sex | RxA |
|---|---|---|---|---|
| **All main effects**<br>~Regime+Age+Sex | **59 (3873)**<br>C>S: 53 (2076)<br>S>C: 6 (1797) | **72 (6385)**<br>Young>Old: 2 (4011)<br>Old>Young: 70 (2374) | **112 (12544)**<br>M>F: 110 (7372)<br>F>M: 2 (5172) | |
| **Males**<br>~Regime+Age | **41 (2858)**<br>C>S: 36 (1690)<br>S>C: 5 (1168) | **108 (7071)**<br>Young>Old: 1 (3556)<br>Old>Young: 107 (3515) | | |
| **Females**<br>~Regime+Age | **27 (2416)**<br>C>S: 23 (1215)<br>S>C: 4 (1201) | **15 (2021)**<br>Young>Old: 5 (1091)<br>Old>Young: 10 (930) | | |
| **Males**<br>~Regime+Age+RxA | | | | **0 (1396)** |
| **Females**<br>~Regime+Age+RxA | | | | **28 (1159)** |
| **Shared[a]** | 19 (1038) | 13 (1479) | | **0 (262)** |
| **Same Direction[a]** | 19 (1002) | 10 (1356) | | |

[a] Number of significant TEs and genes shared between the male and female models (Shared) and number that show the same $\log_2$ FC directionality between levels (Same Direction).

**Table S13 (Excel File). Differential expression analysis output.** Raw RNA-seq reads of Carnes2015 were mapped against a repeat-masked *D. melanogaster* reference genome and TE consensus sequence library using STAR. We then analyzed differential expression between sexes (males, M vs females, F), ages (young vs old), and regimes (selected vs control) using read counts of genes and TE families in DESeq2. The different sheets show the DESeq2 outputs for the different models we used (also see **Table S12**). For the interaction model, we chose young and control as reference levels of age and regime.

**Table S14:** $Log_2$ fold change of 19 TE families differentially expressed across regimes in males and females at an adjusted P-value < 0.05. Positive values denote higher expression in selected, negative values upregulation in controls.

| **TE family** | **Subclass** | **$log_2$ S/C males** | **$log_2$ S/C females** | **Same Direction[a]** |
|---|---|---|---|---|
| *TART-B* | non-LTR | 0.76 | 1.01 | yes |
| *TART-A* | non-LTR | 0.62 | 0.71 | yes |
| *TAHRE* | non-LTR | 0.49 | 0.48 | yes |
| *Quasimodo* | LTR | -0.19 | -0.45 | yes |
| *G5A* | non-LTR | -0.27 | -0.27 | yes |
| *invader2* | LTR | -0.29 | -0.38 | yes |
| *Doc3-element* | non-LTR | -0.29 | -0.31 | yes |
| *17.6* | LTR | -0.34 | -0.36 | yes |
| *Porto1* | non-LTR | -0.47 | -0.39 | yes |
| *gypsy11* | LTR | -0.49 | -0.72 | yes |
| *Rt1b* | non-LTR | -0.52 | -0.39 | yes |
| *mdg1* | LTR | -0.57 | -1.16 | yes |
| *flea* | LTR | -0.58 | -1.04 | yes |
| *gypsy7* | LTR | -0.61 | -0.64 | yes |
| *G5* | non-LTR | -0.69 | -0.44 | yes |
| *ZAM* | LTR | -0.73 | -0.90 | yes |
| *BS* | non-LTR | -0.98 | -0.49 | yes |
| *blood* | LTR | -1.42 | -1.81 | yes |
| *copia* | LTR | -2.48 | -2.29 | yes |

[a]Same Direction denotes TEs that show the same sign of fold change in males and females.

**Table S15:** $Log_2$ fold change of 13 TE families differentially expressed across regimes in males and females at an adjusted P-value < 0.05. Positive values denote higher expression in old, negative values upregulation in young flies.

| TE family | Subclass | log2 Old/Young males | log2 Old/Young females | Same Direction[a] |
|---|---|---|---|---|
| *copia* | LTR | 2.01 | 1.61 | yes |
| *R1A1-element* | non-LTR | 1.92 | 1.22 | yes |
| *springer* | LTR | 1.87 | 1.75 | yes |
| *G-element* | non-LTR | 1.63 | -0.43 | no |
| *R2-element* | non-LTR | 1.52 | 1.29 | yes |
| *Burdock* | LTR | 0.99 | 0.72 | yes |
| *G2* | non-LTR | 0.98 | -0.30 | no |
| *opus* | LTR | 0.89 | 0.64 | yes |
| *mdg1* | LTR | 0.84 | 1.00 | yes |
| *jockey* | non-LTR | 0.83 | 0.73 | yes |
| *rooA* | LTR | 0.77 | 0.13 | yes |
| *Tabor* | LTR | 0.23 | -0.46 | no |
| *297* | LTR | -0.26 | -0.88 | yes |

[a]Same Direction denotes TEs that show the same sign of fold change in males and females.

**Table S16.** Spearman's correlation of normalized expression read counts of TEs to the average number of insertions per sample group, and for averaged expression read counts across all samples.

| Sex | Age | Regime | $\rho^a$ |
|---|---|---|---|
| females | young | control | 0.71 |
| females | young | selected | 0.74 |
| females | old | control | 0.68 |
| females | old | selected | 0.71 |
| Average Expression | | | 0.72 |

[a]All correlation coefficients were significantly different from 0 after Bonferroni correction at $\alpha$ = 0.01.

**Table S17.** TEs significantly different in genomic insertion numbers (Ins.) and in expression (Expr.) between female selected and control populations of Carnes2015. Statistics for TE insertion differences are shown in **Table S2** and differential expression analysis in **Table S13**.

| TE | Ins. (S) | Ins. (C) | δIns. | $log_2$ Ins. | $log_2$ Expr. | Same Direction[a] |
|---|---|---|---|---|---|---|
| *17.6* | 33.12 | 41.77 | -8.65 | -0.34 | -0.36 | yes |
| *1731* | 15.04 | 11.02 | 4.02 | 0.45 | 0.46 | yes |
| *Bari1* | 73.19 | 53.22 | 19.98 | 0.46 | -0.52 | no |
| *blood* | 26.46 | 31.52 | -5.06 | -0.25 | -1.81 | yes |
| *BS* | 11.47 | 10.12 | 1.36 | 0.18 | -0.49 | no |
| *copia* | 55.70 | 107.46 | -51.76 | -0.95 | -2.29 | yes |
| *Dm88* | 19.70 | 15.58 | 4.12 | 0.34 | -0.46 | no |
| *Doc3-element* | 26.52 | 27.70 | -1.18 | -0.06 | -0.31 | yes |
| *G4* | 13.32 | 10.08 | 3.24 | 0.40 | -0.31 | no |
| *G5* | 6.83 | 5.59 | 1.25 | 0.29 | -0.44 | no |
| *G5A* | 6.88 | 4.63 | 2.25 | 0.57 | -0.27 | no |
| *gypsy7* | 7.45 | 5.58 | 1.88 | 0.42 | -0.64 | no |
| *invader2* | 30.92 | 32.49 | -1.57 | -0.07 | -0.38 | yes |
| *jockey* | 34.70 | 29.01 | 5.69 | 0.26 | -0.51 | no |
| *mdg1* | 40.41 | 31.73 | 8.69 | 0.35 | -1.16 | no |
| *Porto1* | 14.46 | 18.73 | -4.27 | -0.37 | -0.39 | yes |
| *Quasimodo* | 45.96 | 34.90 | 11.05 | 0.40 | -0.45 | no |
| *R1A1-element* | 122.48 | 154.65 | -32.17 | -0.34 | -1.32 | yes |
| *rooA* | 11.66 | 6.43 | 5.23 | 0.86 | -0.17 | no |
| *Rt1b* | 24.91 | 21.62 | 3.29 | 0.20 | -0.39 | no |
| *TAHRE* | 11.03 | 10.35 | 0.68 | 0.09 | 0.48 | yes |
| *Transpac* | 29.91 | 26.60 | 3.31 | 0.17 | -0.68 | no |
| *ZAM* | 4.66 | 7.11 | -2.45 | -0.61 | -0.90 | yes |

[a]Same Direction denotes significant TEs with the same sign for δIns. and $log_2$ expression differences.

**Table S18 (Excel File). Differences in average expression per TE insertion.** We divided filtered RNA-seq read by the number of TE insertions in each population (see **Table S2**). We then analyzed differences in average expression per TE insertion in females between ages (young vs old), and regimes (selected vs control) using DESeq2. The different sheets show a summary of significant TE families and genes at an adjusted P-value < 0.05, and the outputs for the different models. For the interaction model (RxA, Regime x Age), we chose young and control as reference levels of age and regime. Sample sizes varied between models and factors dependent on auto-filtering by DESeq2, and ranged from 120 to 123 TEs and 10,850 to 12,911 genes. The median sample size across all factors was n = 122 for TEs and n = 11,622 for genes.

**Table S19 (Excel File). Genomic and transcriptomic candidate genes implicated in TE regulation.** Genes likely involved in TE regulation and transposition (i.e. chromatin-structure, piRNA pathway, and transposition-associated genes) were obtained from the literature and the gene ontology consortium and broadly classified into three types (epigenetic, piRNA, transposition). We compared the list of TE regulation genes to the reported candidate gene lists from Carnes2015, Fabian2018, Hoedjes2019 and Remolina2012. We further re-analyzed RNA-seq data from Carnes2015 (whole flies, males and females, 2 ages) and a microarray experiment from Remolina2012 (female heads and abdomens, 5 ages) to investigate if TE regulation genes vary across age and selection regimes. "S>C" denotes that the gene is higher expressed in selected populations and "C>S" a higher expression in controls. "Old>Young" means higher expression in old flies and "Young>Old" higher expression young flies. "Increasing" or "decreasing" denotes that the slope of the age effect was positive (enhanced expression with age) or negative (decreased expression with age), respectively. Significance was defined by an FDR < 0.05 unless otherwise indicated. Annotation of FlyBase IDs (FBgn) and gene names are based on the *D. melanogaster* reference v6.27.

**Table S20.** Programs and options used for the creation of files for analyses.

| Process / analysis | Program | Options |
|---|---|---|
| ***Genome-wide TE abundance*** | | |
| TE copy numbers from raw fastq files | DeviaTE | 1) --read_type phred+33 (phred+64 for Fabian2018), --min_read_len 50, --quality_threshold 18, --hq_threshold 20, --min_alignment_len 30, --threads 20, --families ALL, --single_copy_genes Dmel_rpl32,Dmel_RpII140,Dmel_Act5C,Dmel_piwi,Dmel_p53<br>2) Summed high quality coverage ('hq_cov') and physical coverage ('phys_cov') columns for each consensus sequence position to obtain *NormalizedCoverage* (Fig.S1). Due to sequence similarity within and between TEs, repeated most analyses without filtering for mapping quality using values from the 'cov' column instead of 'hq_cov'. |
| ***Genomic TE locations and activity*** | | |
| Trimming | cutadapt (v.2.0) | --minimum-length 50 -q 18 (also --quality-base=64 for Fabian2018) |
| Repeat-masking of reference gnome (v.6.27) | RepeatMasker (v.4.0.9) with RMBlast (v.2.9.0) | -gccalc -s -cutoff 200 -no_is -nolow -norna -gff -u -pa 18 |
| Mapping | bwa bwasw (v.0.7.17) | |
| Genomic TE insertion location | PoPoolationTE2 | *1) se2pe*<br>2) *ppileup* --map-qual 15<br>3) *identifySignatures* --signature-window fix500 --min-valley fix150 --mode joint; due to differences in the number of replicate populations we chose --min-count 3 for Remolina2012 and Fabian2018, --min-count 5 for Carnes2015, or --min-count 12 for Hoedjes2019<br>4) frequency<br>5) filterSignatures --max-otherte-count 2 --max-structvar-count 2<br>6) pairupSignatures --min-distance -200 --max-distance 300 |
| ***Genome-wide nucleotide diversity*** | | |
| Mapping | bwa *mem* (v.0.7.17) | |
| Bam files | samtools (v.1.9) | 1) samtools view -Sb<br>2) samtools view sort |
| Duplicate filtering | PicardTools (v.2.18.27) | MarkDuplicates with REMOVE_DUPLICATES=true. |
| Merging lanes in Remolina2012 | samtools *merge* | |
| mpileup file | samtools mpileup | -q 20 --rf 0x2 --ff 0x4 --ff 0x8 -f (and -6 for Fabian2018) |
| Nucleotide diversity $\pi$ and Watterson's $\theta$ | Popoolation | Variance-sliding.pl --min-count 2 --min-coverage 5 --window-size 100000 --step-size 100000 --pool-size 100 (or 250 for Hoedjes2019) --measure pi (or theta) --fastq-type sanger --min-covered-fraction 0.6<br>We set the maximum coverage threshold (--max-coverage) 2x of the average genome coverage of each population. The range of cut-offs |

| | | |
|---|---|---|
| | | were: 20-74 in Carnes2015, 286-355 in Fabian2018, 67-107 in Remolina2012 and 166-228 in Hoedjes2019. |
| ***RNA-seq analysis*** | | |
| Trimming | cutadapt (v.2.0) | -a AGATCGGAAGAGC --minimum-length 75 -q 20 |
| Mapping | STAR (v2.7.0e) | --alignIntronMin 23 --alignIntronMax 268107 -–outFilterMultimapNmax 10 --outReadsUnmapped FastX --outSAMstrandField intronMotif --outSAMtype BAM SortedByCoordinate |
| Read Counts | featureCounts in subread 1.6.3 | -t exon -g gene_id --extraAttributes gene_symbol |
| Differential expression | DESeq2 in R | |

#### SUPPLEMENTARY FIGURES

**Figure S1. Overview of three approaches used to test differences in TE abundance by the example of the telomeric TE *HeT-A*.** (A) We used DeviaTE (Weilguny and Kofler 2019) to quantify the number of insertions for each TE family. The tool quantifies the sequencing depth across the consensus positions within a TE family considering the physical coverage of split reads spanning deletions. To obtain copy number estimates, DeviaTE uses the coverage of every TE family and normalizes it by the depth of the same five single-copy genes (see **Fig. S2**), which results in normalized coverage values (NormCov) for each position. In our first approach (Approach #1), we use NormCov estimates of all consensus sequence positions assuming they are independent measures to analyze studies separately. We fit a model for each TE family with the NormCov values as dependent variable to test significant differences between selection ("Sel", late-breeding, long-lived) and control ("Cont", earlier breeding, shorter lifespan) regimes, and Bonferroni-corrected P-values for multiple testing (at $\alpha = 0.01$). The resulting statistic is not conservative and comparable to just considering average differences between control and selected populations but allowed us to identify TE families differing between regimes within a study despite the small number of replicate populations. In the line plot, solid lines and shaded areas indicate the average and variability across populations within regime of Hoedjes2019, respectively. The boxplot shows the overall difference in NormCov values between control and selected populations. (B) To obtain single mean insertion estimates per TE family and population, we averaged the NormCov values across all consensus sequence positions within a TE family. The mean insertion estimates were used for all the downstream analysis. We also obtained the total genomic TE content by summing up all mean insertion counts across detected TEs within a population. Because the total genomic TE content was significantly different between studies (see **Fig. 1D**, **Table S6),** we divided the insertion estimates of all TE families by the total insertions in the corresponding population and used these proportions for approach #2. (C) For Approach #2, we arcsine square root transformed these insertion proportions (PropIns) and test differences in abundance across regimes for each TE family combining all studies, followed by FDR correction (<0.05) to account for multiple testing. (D) In our most conservative approach (Approach #3), we identified TE families that had consistently higher mean insertion counts across all selected populations compared to controls and vice versa.

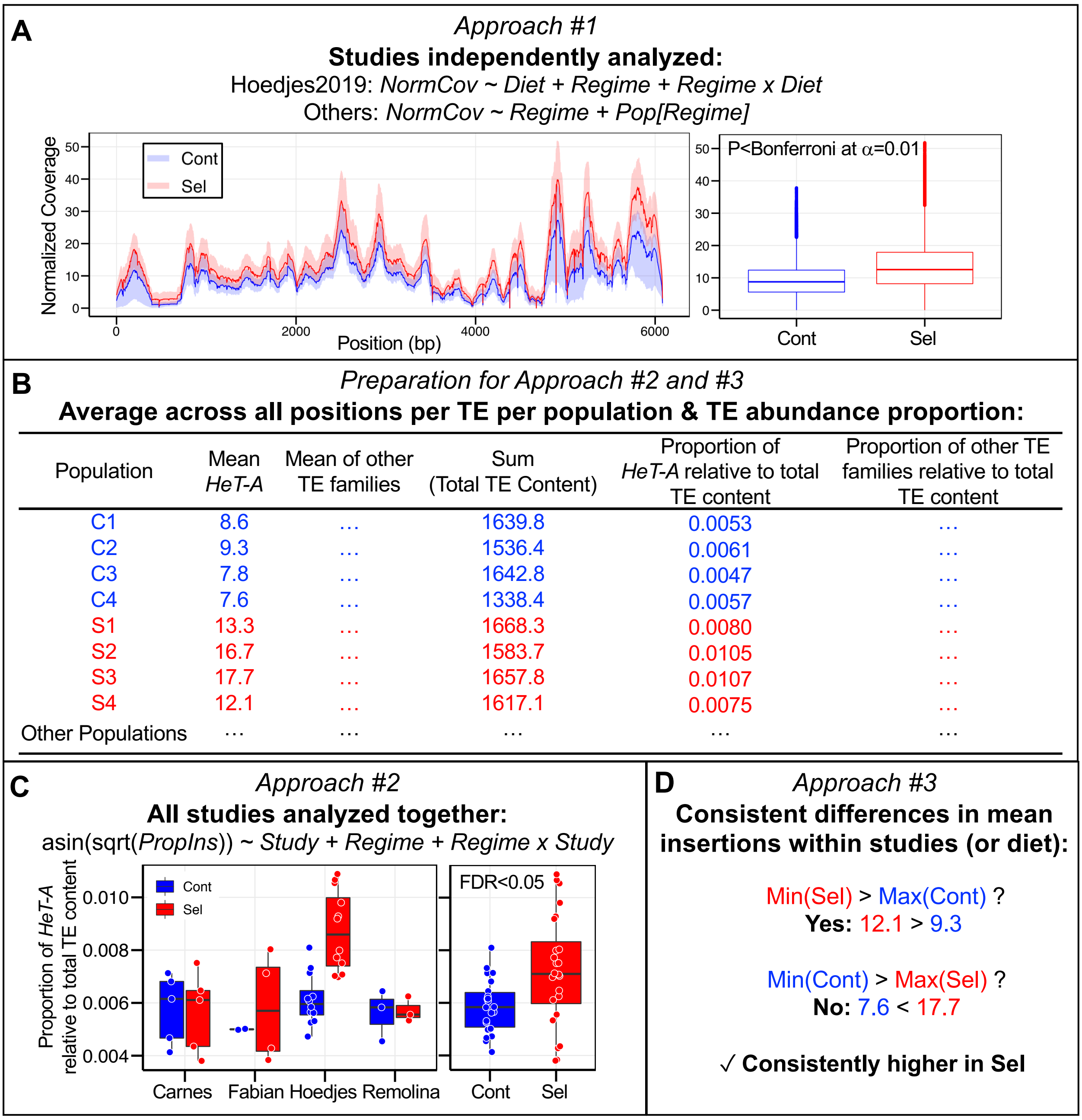
A
Approach #1
Studies independently analyzed:
Hoedjes2019: NormCov ~ Diet + Regime + Regime x Diet
Others: NormCov ~ Regime + Pop[Regime]
Normalized Coverage
Cont
Sel
Position (bp)
P<Bonferroni at α=0.01
Cont
Sel
B
Preparation for Approach #2 and #3
Average across all positions per TE per population & TE abundance proportion:
Population | Mean HeT-A | Mean of other TE families | Sum (Total TE Content) | Proportion of HeT-A relative to total TE content | Proportion of other TE families relative to total TE content
C1 | 8.6 | … | 1639.8 | 0.0053 | …
C2 | 9.3 | … | 1536.4 | 0.0061 | …
C3 | 7.8 | … | 1642.8 | 0.0047 | …
C4 | 7.6 | … | 1338.4 | 0.0057 | …
S1 | 13.3 | … | 1668.3 | 0.0080 | …
S2 | 16.7 | … | 1583.7 | 0.0105 | …
S3 | 17.7 | … | 1657.8 | 0.0107 | …
S4 | 12.1 | … | 1617.1 | 0.0075 | …
Other Populations | … | … | … | … | …
C
Approach #2
All studies analyzed together:
asin(sqrt(PropIns)) ~ Study + Regime + Regime x Study
Proportion of HeT-A relative to total TE content
Cont
Sel
FDR<0.05
Carnes
Fabian
Hoedjes
Remolina
Cont
Sel
D
Approach #3
Consistent differences in mean insertions within studies (or diet):
Min(Sel) > Max(Cont) ?
Yes: 12.1 > 9.3
Min(Cont) > Max(Sel) ?
No: 7.6 < 17.7
✓ Consistently higher in Sel

**Figure S2. Average coverage of single-copy genes.** The coverage of five single-copy genes used by DeviaTE to normalize TE family sequencing depth are shown for each replicate population and study. Populations are labelled according to regime and their published names. Cont, Control; Sel, Selection regime. For Hoedjes2019: L, low sugar/protein diet, M, medium sugar/protein; H; high sugar/protein; E, early reproduction; P, postponed reproduction.

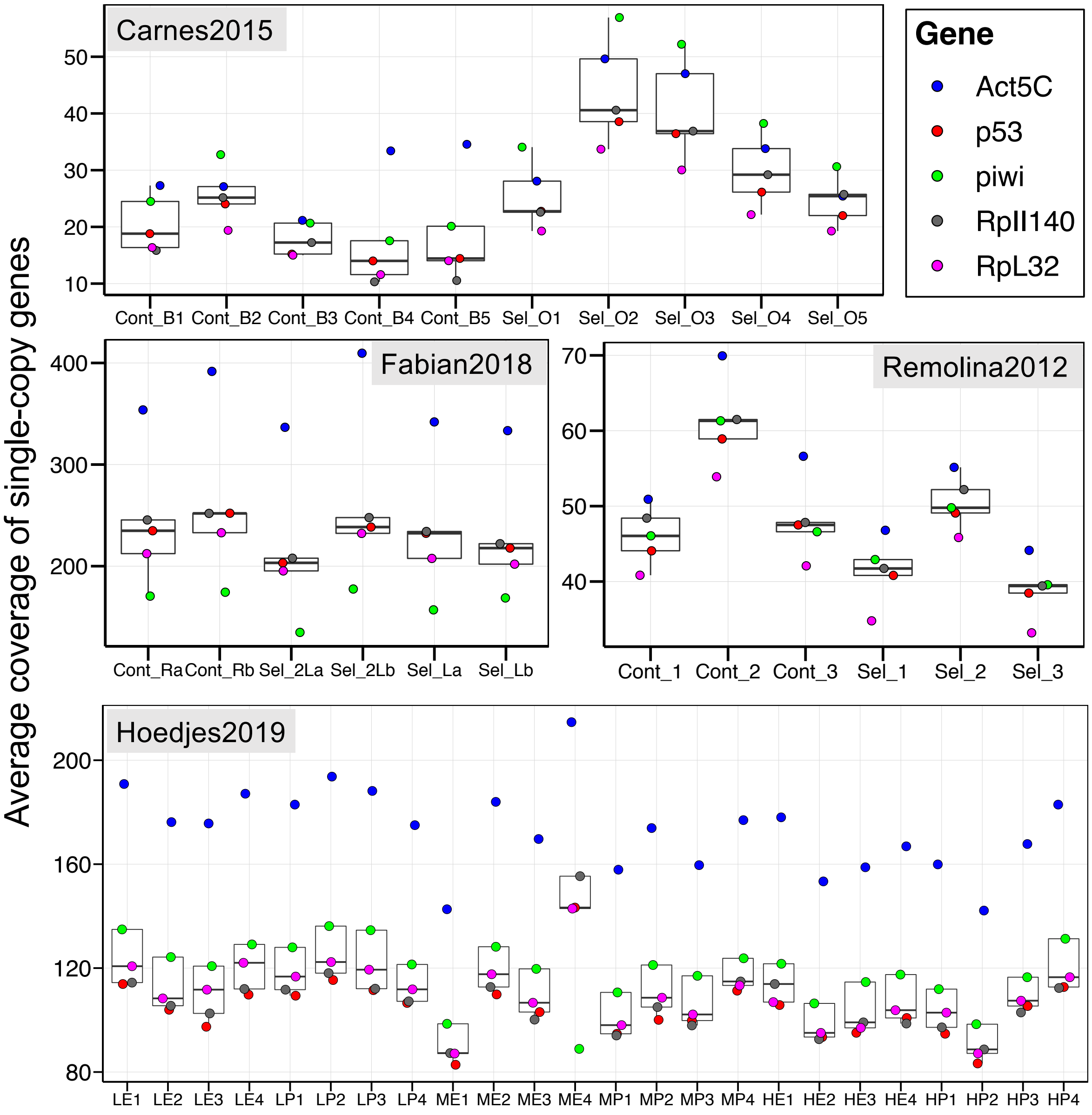

**Figure S3. Filtering TE families by normalized coverage and minimum number of covered positions.** Each point represents one of 179 TE families included in DeviaTE. The Y-axis shows the proportion of TE family consensus sequence positions with normalized coverage (NormCov) >=0.5. The solid red line denotes our chosen cut-off, i.e. a minimum NormCov of 0.5 across at least 80% of the positions within a TE family. Thus, TE families without any mapped reads (i.e. 24-29 across studies), and with very low and/or very few insertion estimates were excluded from the downstream analysis. Dependent on the study, this resulted in 110-115 retained TE families. 103 TE families were shared across all studies after filtering. Results were qualitatively similar when we did not filter for coverage or mapping quality (not shown). Please also see **Fig. S1A** and Materials and Methods.

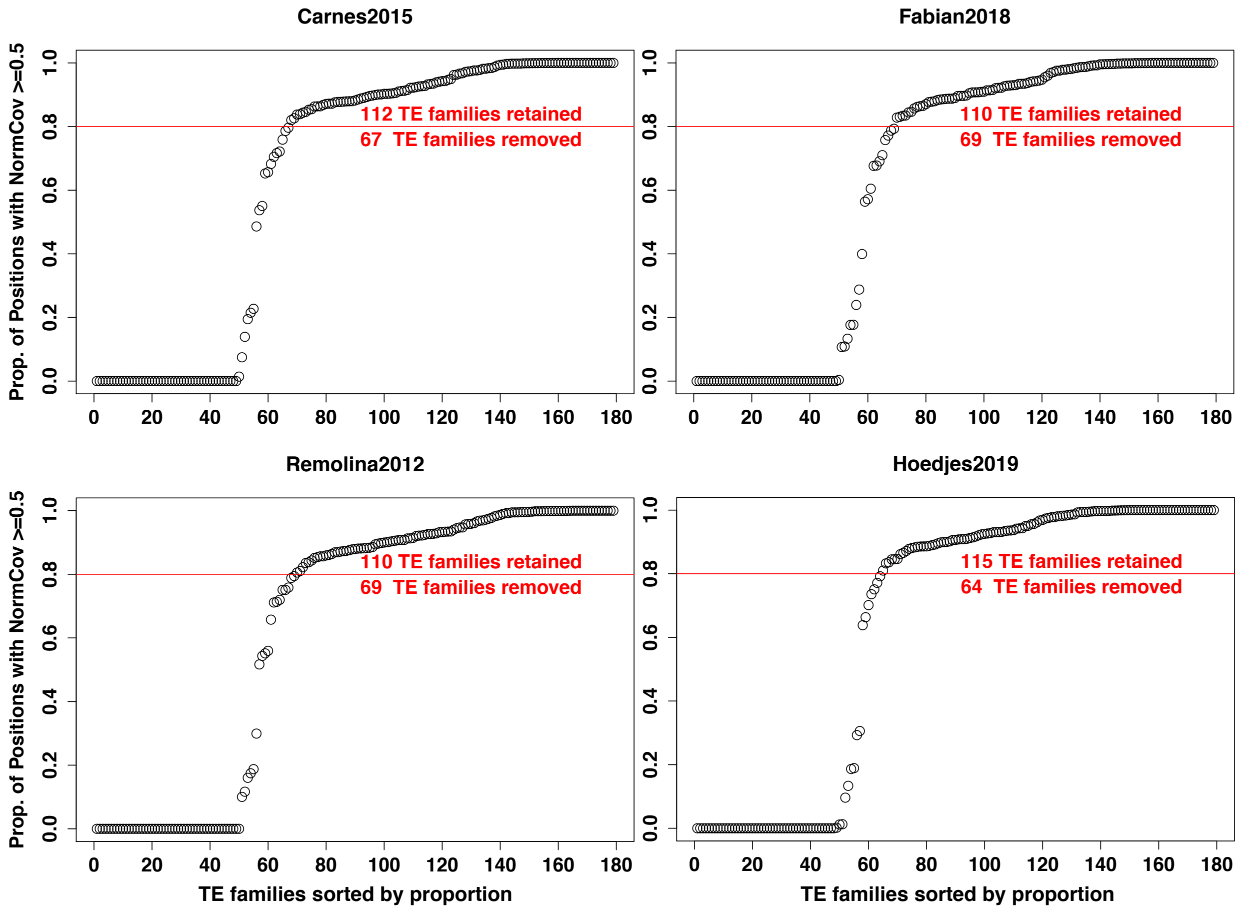

**Figure S4. Difference in TE abundance between selected and control populations in different diet conditions of Hoedjes2019.** Populations were selected on three different larval diets ranging from low to high sugar/protein content. Upper row: δinsertions (Y-axis) were obtained by subtracting the average genomic insertions of the controls from the selected populations. Bottom row: $log_2$ fold change between selected and control populations. The dashed line indicates no difference between regimes. >0 denote TEs with a larger abundance in selected populations, while <0 TEs with more insertions in controls. TE subclasses are given in different colors. Selected populations had more genomic insertions than controls for most TE families when maintained on the low and medium larval food environment, while the high diet showed the opposite pattern (also see **Table S2** and **Table S3**).

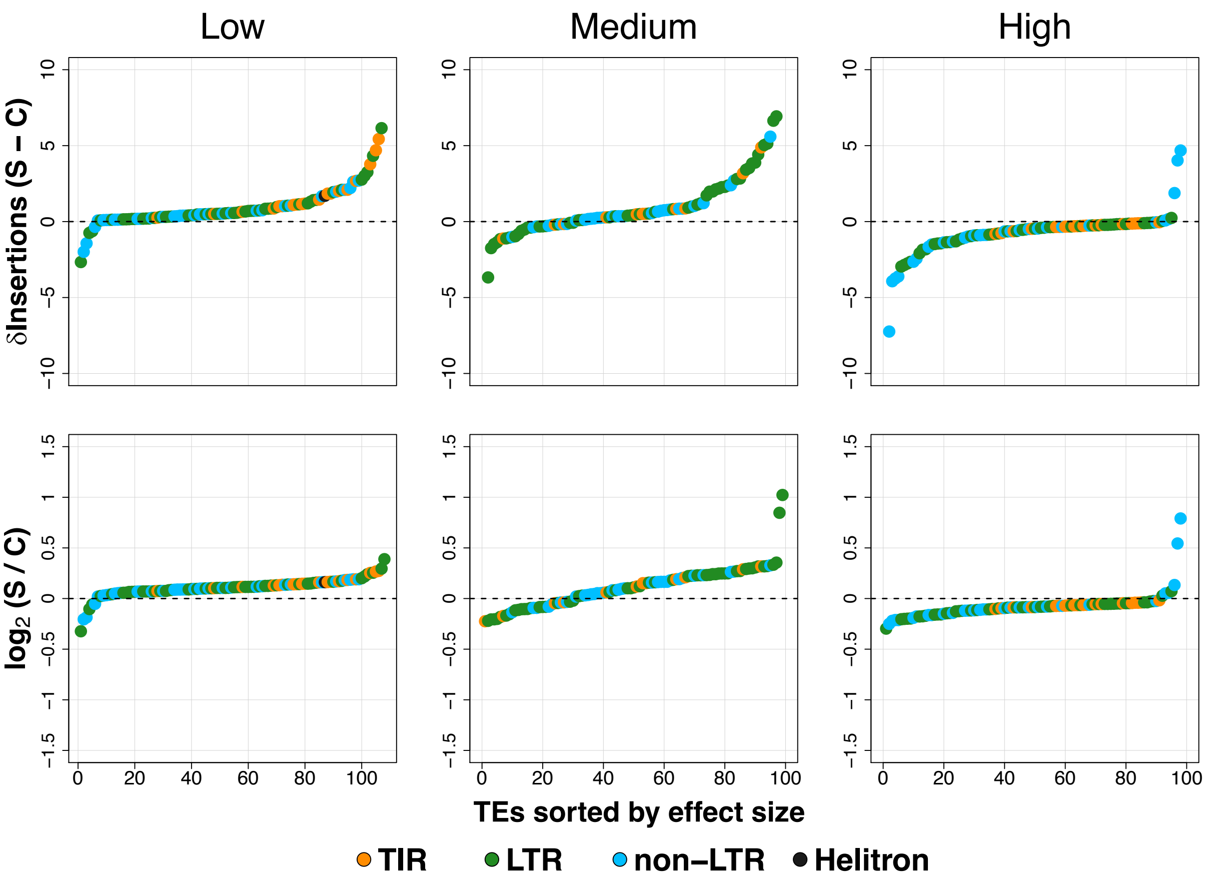

**Figure S5. Difference in TE abundance between regimes.** (A) δInsertion (Y-axis) were obtained by subtracting the average genomic insertions of the controls ("C") from the selected populations ("S"). The dashed line indicates no difference between regimes. >0 denote TE families with a larger abundance in selected populations ("S>C"), while <0 TEs with more insertions in controls ("C>S"). TE subclasses are given in different colors. Selected flies had more genomic insertions than controls for most TE families (also see **Table 1**). (B) Difference in magnitude of absolute δInsertion change between C>S and S>C TE groups. Significant difference between TE groups was determined using t-tests independently for each study. * P < 0.05; ** P < 0.01; ns, not significant.

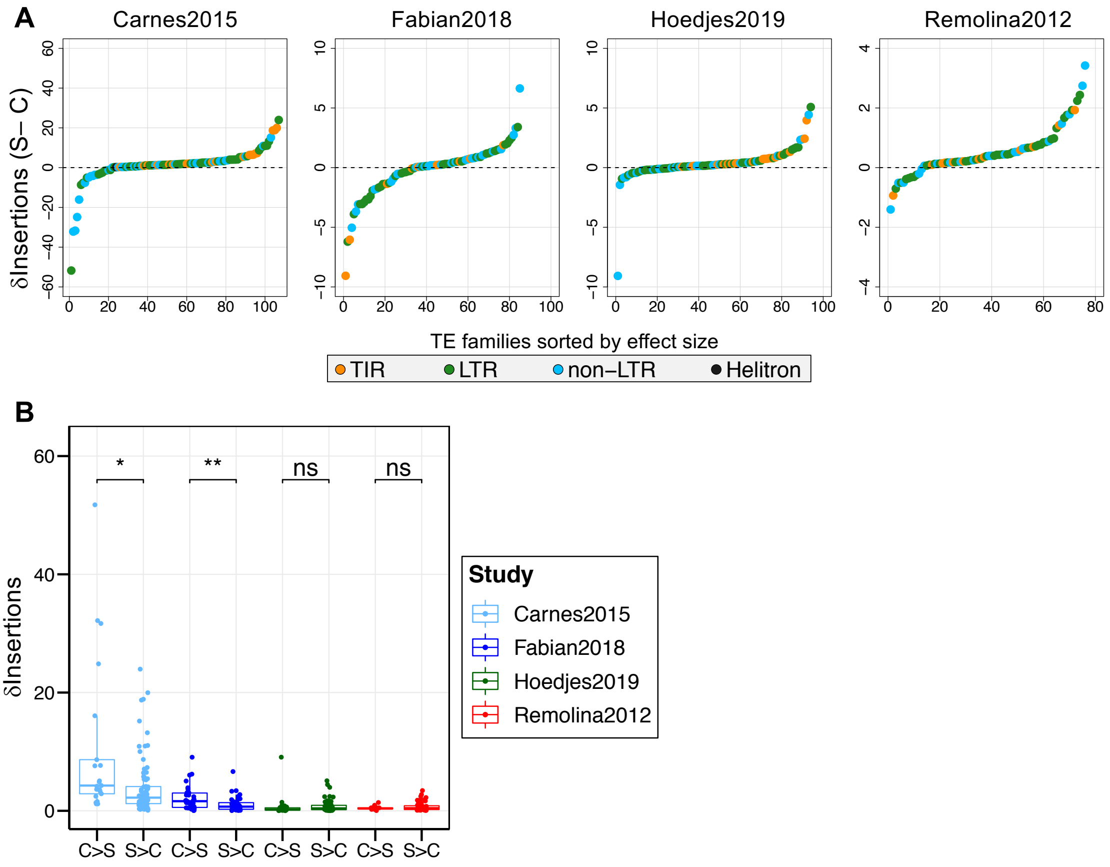

**Figure S6. Number of genomic insertions of TE subclasses**. From top to bottom: sum of all terminal inverted repeats (TIR), long terminal repeats (LTR), and non-long terminal repeat (non-LTR) TEs in the genomes of all populations. ANOVA models combining all studies were used to test the effects of Study, Regime and the Study x Regime interaction. The effect of Study was significant for all TE subclasses. TIRs and LTRs showed a significant increase in selected relative to control populations. See **Table S6** for a summary of the statistical analysis.

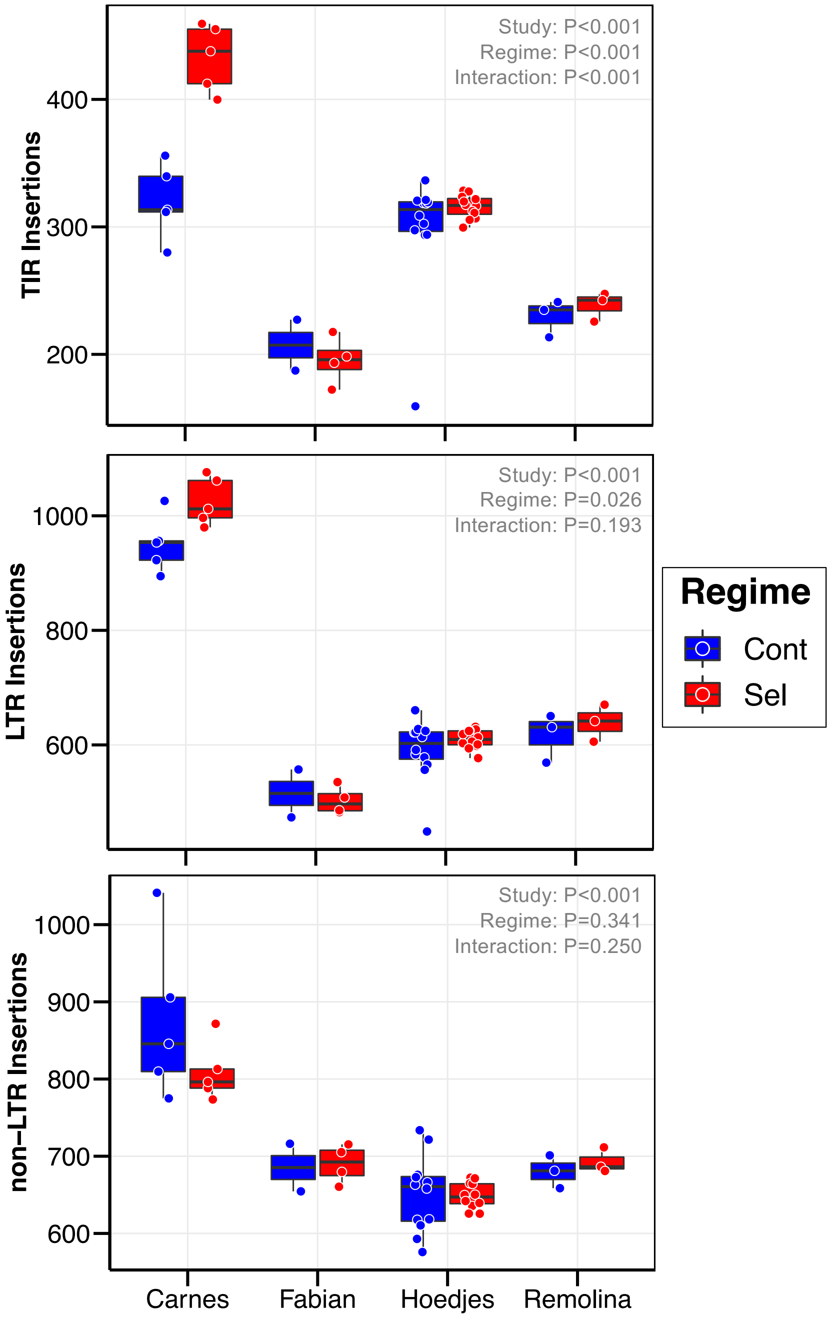

**Figure S7. Spearman's correlation between average population frequencies of experimental evolution studies and South African population.** For every TE family, we calculated average frequency within a population. Spearman's correlation with the South African (SA) population was significant and similar for all populations within three studies (Fabian2018: $\rho = 0.64 – 0.66$; Hoedjes2019: $\rho = 0.59 – 0.63$; Remolina2012: $\rho = 0.57 – 0.61$; all $P < 0.0001$). In Carnes2015, where we only detected a small number of TE insertions possibly due to low coverage, the range of $\rho$ was -0.15 to 0.27 and only one population was significantly correlated (one selected population, $\rho = 0.27$, $P = 0.02$). For simplicity, we here show the relationship between TE frequencies averaged across all populations within a study and the South African population (left panels). Ranks of frequencies are shown in the right panels. TEs with a significant difference in genomic abundance are indicated in red (S>C) or blue (C>S), where grey points denote not significant TEs. The dashed line indicates where TEs would be if there were no differences in frequency. *** $P < 0.0001$; ns, not significant.

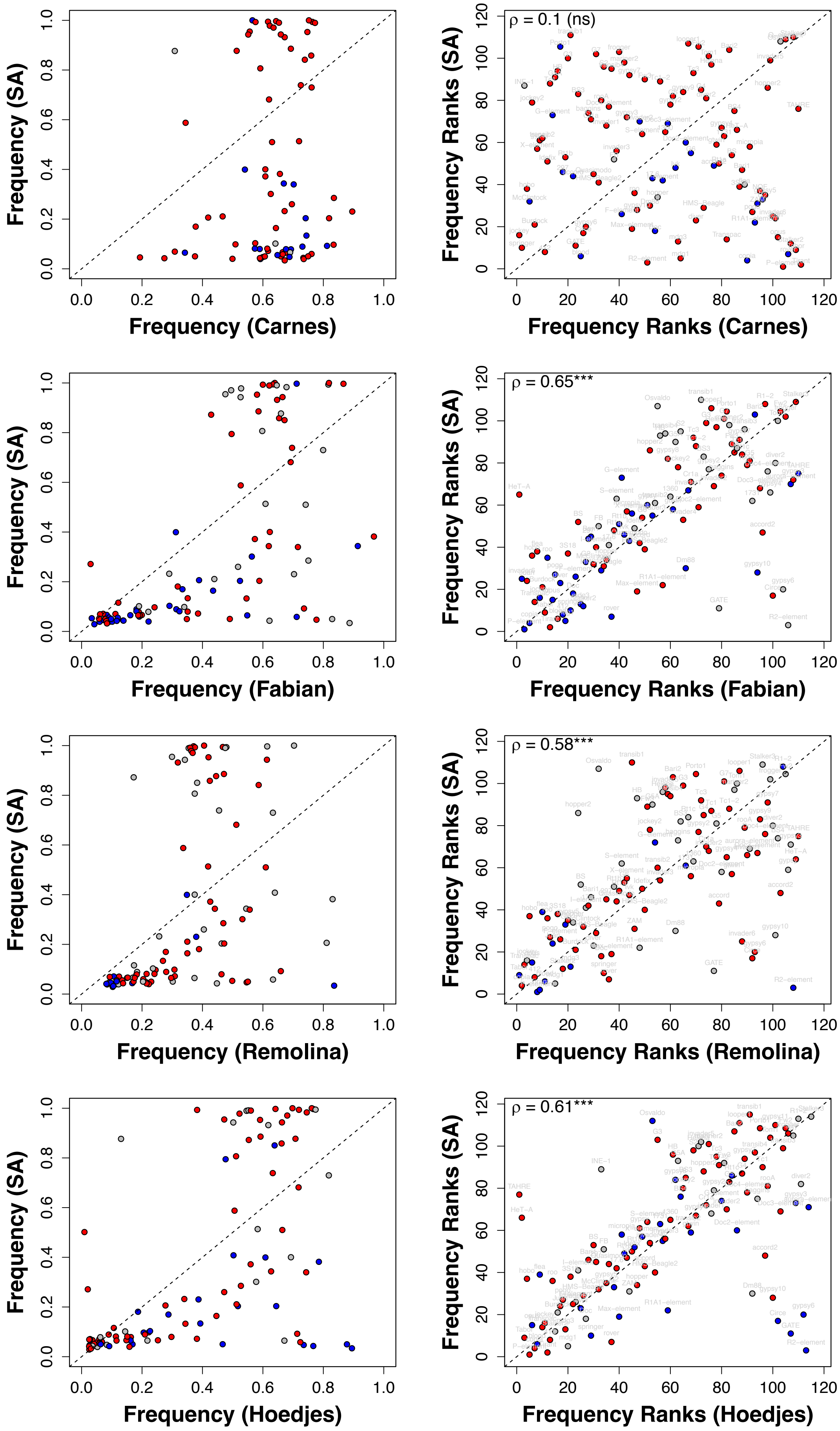

Frequency (SA)
Frequency (Carnes)
Frequency Ranks (SA)
Frequency Ranks (Carnes)
ρ = 0.1 (ns)
Frequency (Fabian)
Frequency Ranks (Fabian)
ρ = 0.65***
Frequency (Remolina)
Frequency Ranks (Remolina)
ρ = 0.58***
Frequency (Hoedjes)
Frequency Ranks (Hoedjes)
ρ = 0.61***

**Figure S8. Spearman's correlation between number of genomic insertions and average population frequencies of the South African population.** The number of genomic TE insertions from populations of the four experimental evolution studies is significantly negative correlated to average TE family frequency from the South African (SA) population (Spearman's $\rho$ range, Carnes2015: $\rho$ = -0.54 to -0.49, Fabian2018: $\rho$ = -0.5 to -0.4; Hoedjes2019: $\rho$ = -0.45 to -0.42; Remolina2012: $\rho$ = -0.51 to -0.5; all $P < 0.0001$). For simplicity, we here show the relationship between number of insertions averaged across all populations within a study and the TE family frequencies in the SA population (left panels). Ranked data are shown in the right panels. TEs with a significant difference in genomic abundance are indicated in red (S>C) or blue (C>S), where grey points denote not significant TEs. *** $P < 0.0001$.

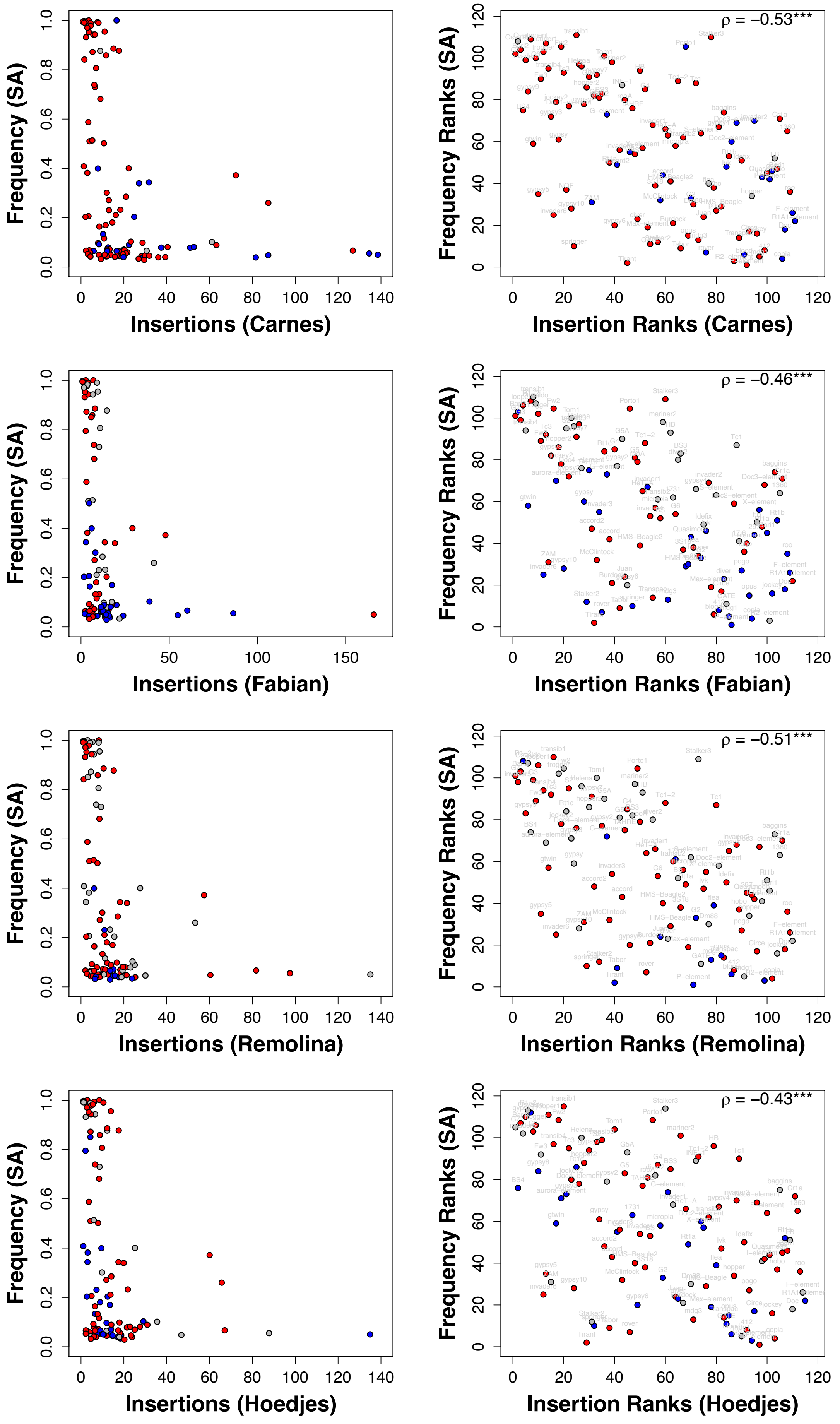

Frequency (SA)
Insertions (Carnes)
Frequency Ranks (SA)
Insertion Ranks (Carnes)
ρ = −0.53***
Frequency (SA)
Insertions (Fabian)
Frequency Ranks (SA)
Insertion Ranks (Fabian)
ρ = −0.46***
Frequency (SA)
Insertions (Remolina)
Frequency Ranks (SA)
Insertion Ranks (Remolina)
ρ = −0.51***
Frequency (SA)
Insertions (Hoedjes)
Frequency Ranks (SA)
Insertion Ranks (Hoedjes)
ρ = −0.43***

**Figure S9. Spearman's correlation between number of genomic insertions and average population frequencies in four experimental evolution studies.** The number of genomic TE insertions is significantly negative correlated to average TE family frequency within populations of Fabian2018, Remolina2012, and Hoedjes2012 (Spearman's $\rho$ range, Fabian2018: $\rho = -0.38$ to $-0.28$; Hoedjes2019: $\rho = -0.41$ to $-0.24$; Remolina2012: $\rho = -0.41$ to $-0.39$; all $P < 0.01$). For Carnes2015, despite 7/10 populations showed negative correlation coefficients, the relationship was only significant in the two populations with the highest and lowest coefficients (Spearman's $\rho$ range, -0.29 to 0.28; for lowest and highest $\rho$, $P = 0.01$; all others not significant). For simplicity, we here show the relationship between number of insertions averaged across all populations and the average TE family frequencies within a study (left panels). Ranked data are shown in the right panels. TEs with a significant difference in genomic abundance are indicated in red (S>C) or blue (C>S), where grey points denote not significant TEs. *** $P < 0.0001$; ** $P < 0.001$; ns, not significant.

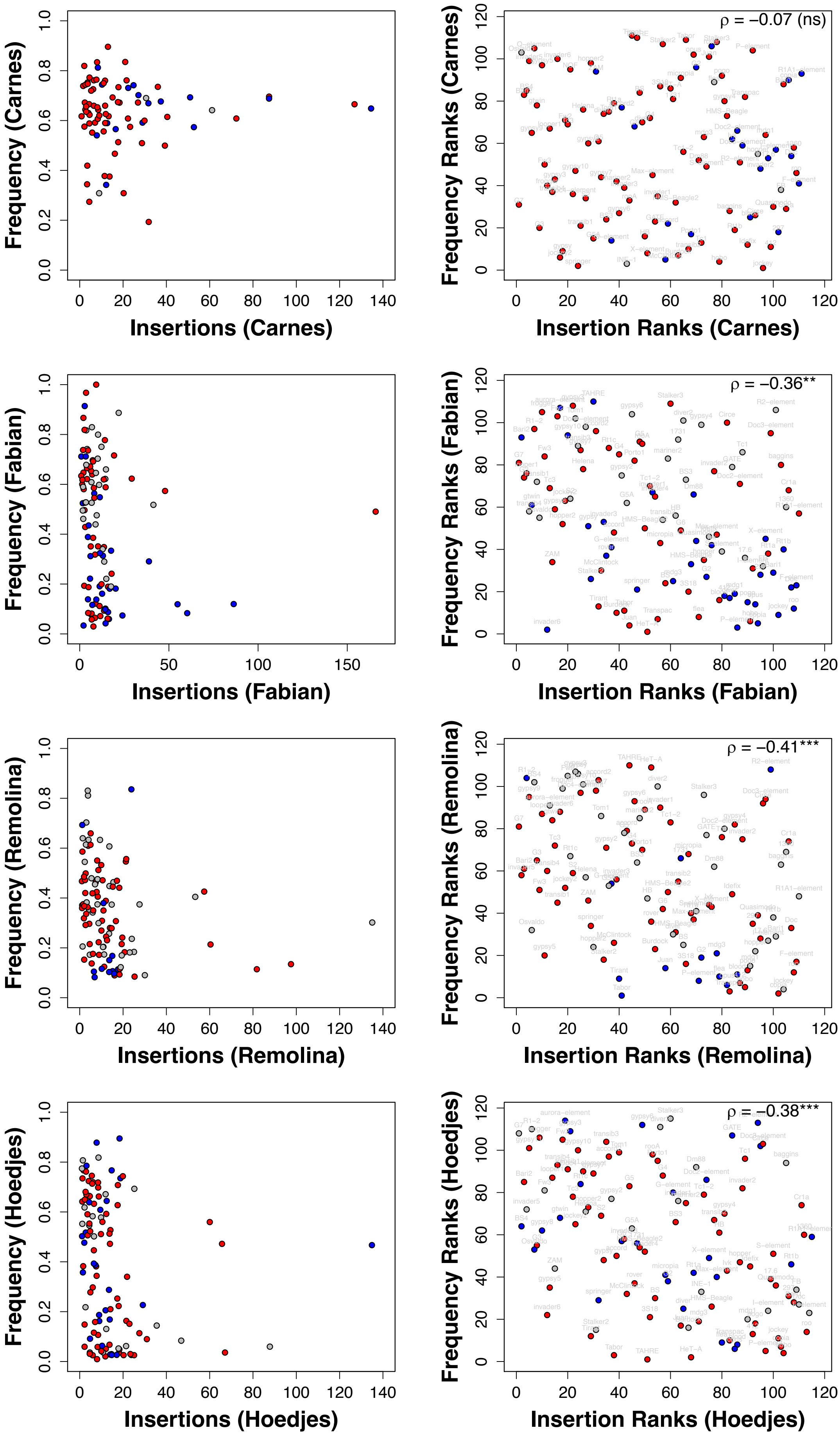
Frequency (Carnes)
Insertions (Carnes)
ρ = −0.07 (ns)
Frequency Ranks (Carnes)
Insertion Ranks (Carnes)
Frequency (Fabian)
Insertions (Fabian)
ρ = −0.36**
Frequency Ranks (Fabian)
Insertion Ranks (Fabian)
Frequency (Remolina)
Insertions (Remolina)
ρ = −0.41***
Frequency Ranks (Remolina)
Insertion Ranks (Remolina)
Frequency (Hoedjes)
Insertions (Hoedjes)
ρ = −0.38***
Frequency Ranks (Hoedjes)
Insertion Ranks (Hoedjes)

**Figure S10. Expected proportions of S>C to C>S TEs under genetic drift.** Histograms show distribution of $\log_2$ relative proportions after 5,000 simulations according to the experimental design of each study (**Table S1**). The red dashed line indicates observed relative proportions when studies were analyzed independently (Table 1). P-value was obtained through dividing the number of simulations with equal or higher values than the observed proportion by the total number of simulations. The black dashed line denotes 0, i.e. equal proportions of S>C and C>S TEs.

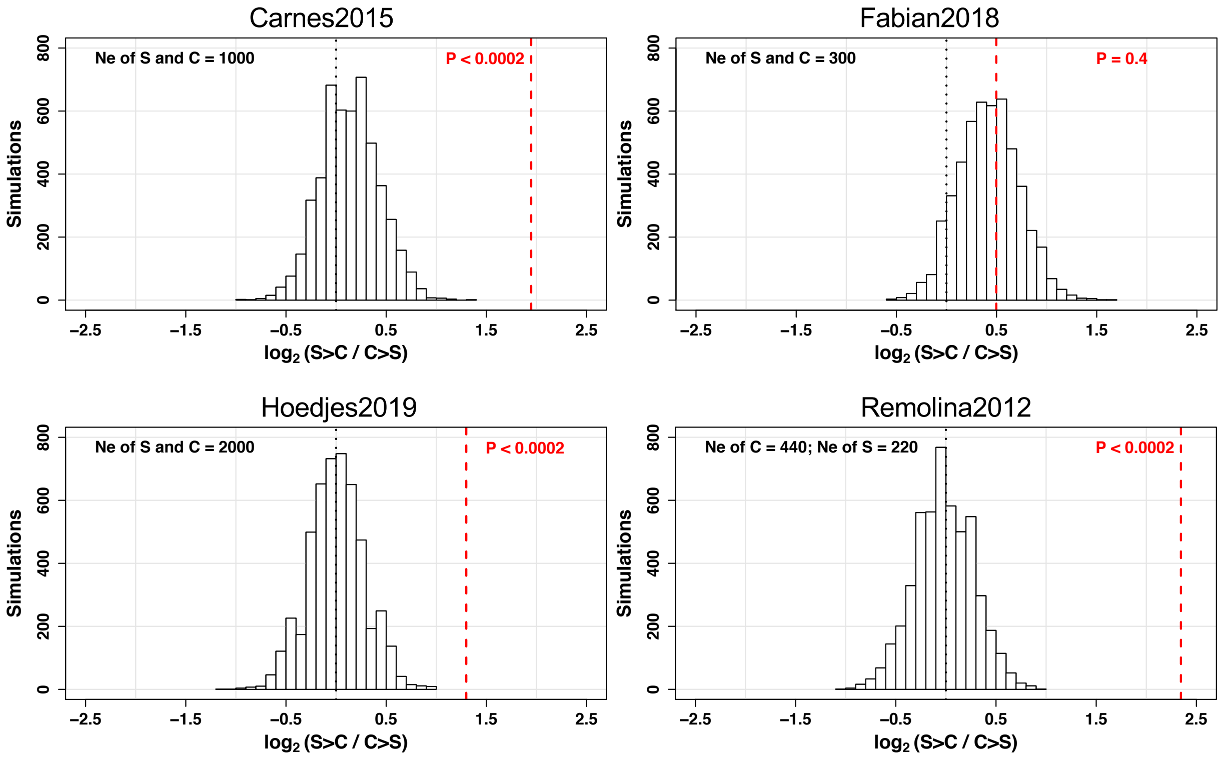

**Figure S11. Differences in TE expression from model with Sex, Age, and Regime.** $Log_2$ fold change in TE family expression of (A) males relative to females, (B) old relative to young age, and (C) selected relative to control populations. 112 TEs significant for sex, 72 for age, and 59 for regime are shown in red, non-significant ones in grey. Error bars are standard errors of the $log_2$ fold change estimate. See **Table S12** for a summary, and **Table S13** for the DESeq2 output.

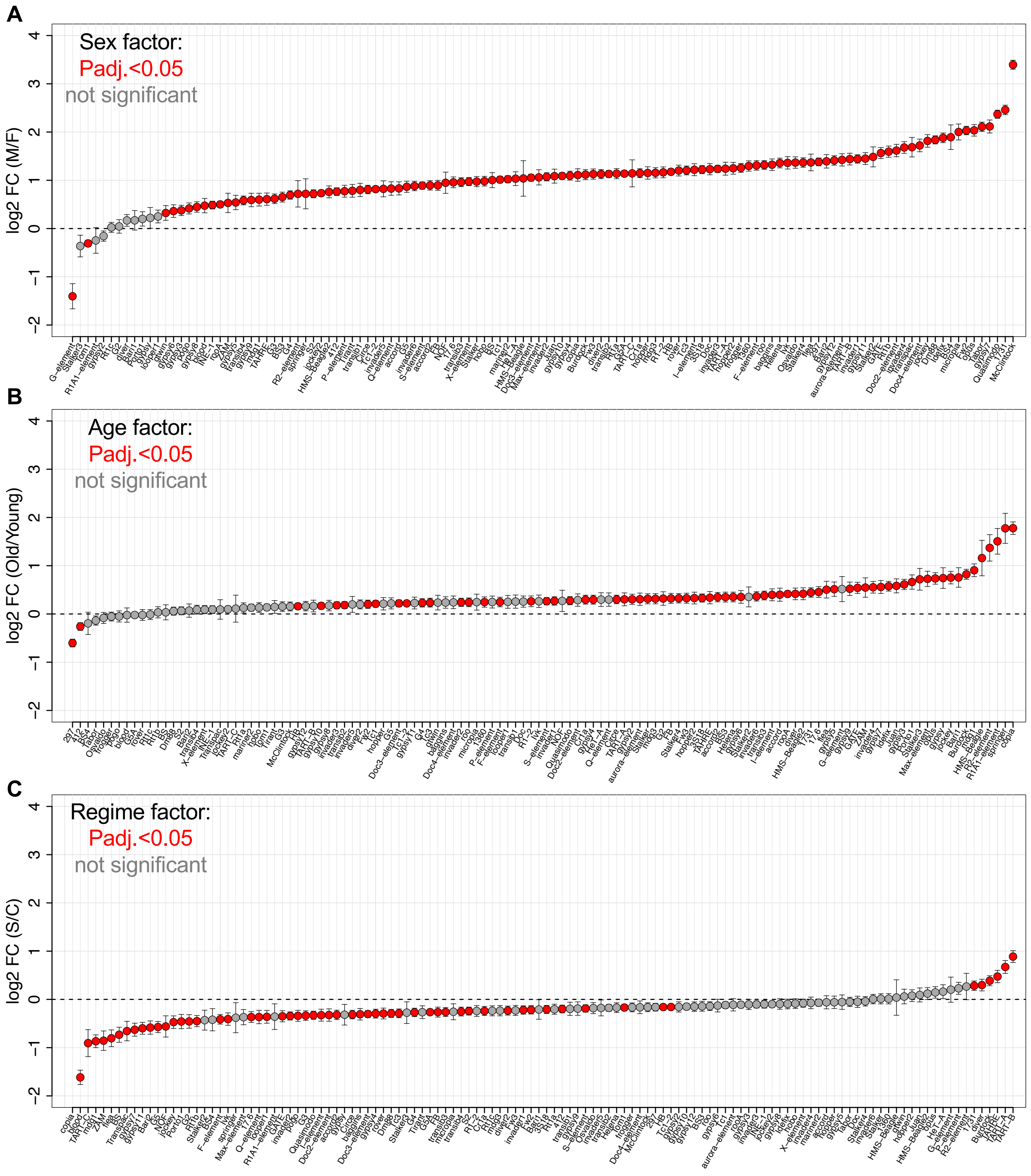

**Figure S12. Magnitude of expression change of TE families across regime and age.** Upper row: Upper, left: boxplots show $log_2$ fold change (Y-axis) of 36 TEs with higher (C>S) and 5 with lower expression (S>C) in control males relative to selected. Upper, right: 23 TEs with higher (C>S) and 4 with lower expression (S>C) in control females relative to selected. Lower, left: 1 TE that decreased and 107 TEs that increased expression with age in males. Lower, right: 5 TEs that decreased and 10 that increased expression with age in females. Statistical differences between groups were analyzed using t-tests.

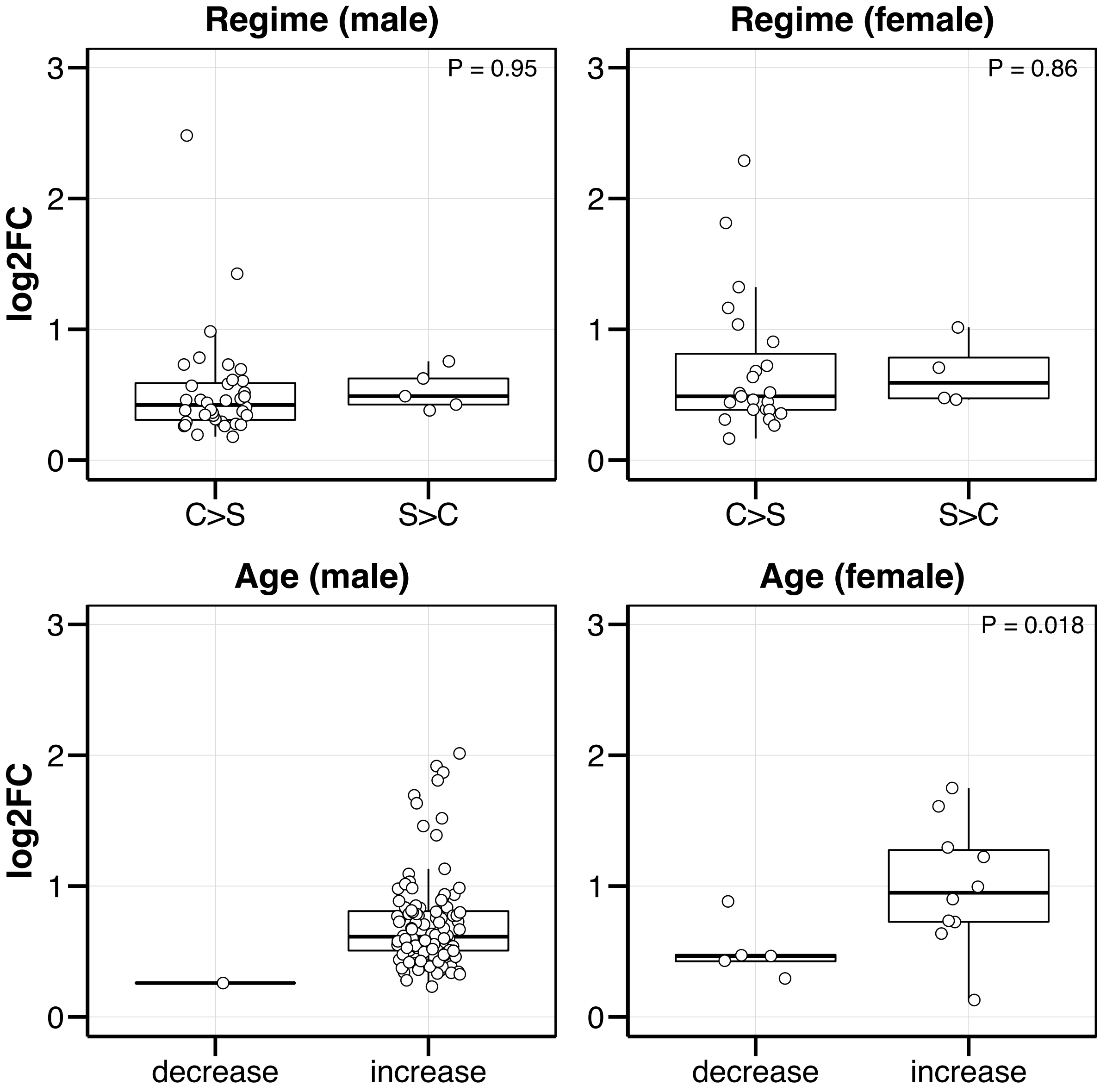

**Figure S13. Example of Regime x Age interaction in females.** Normalized read counts are shown for *Doc3-element* on the Y-axis. Age is on the X-axis. Controls, Cont (blue points); selection lines, Sel (red squares). Among the 28 TEs with a significant interaction, *Doc3-element* had the lowest adjusted P-value.

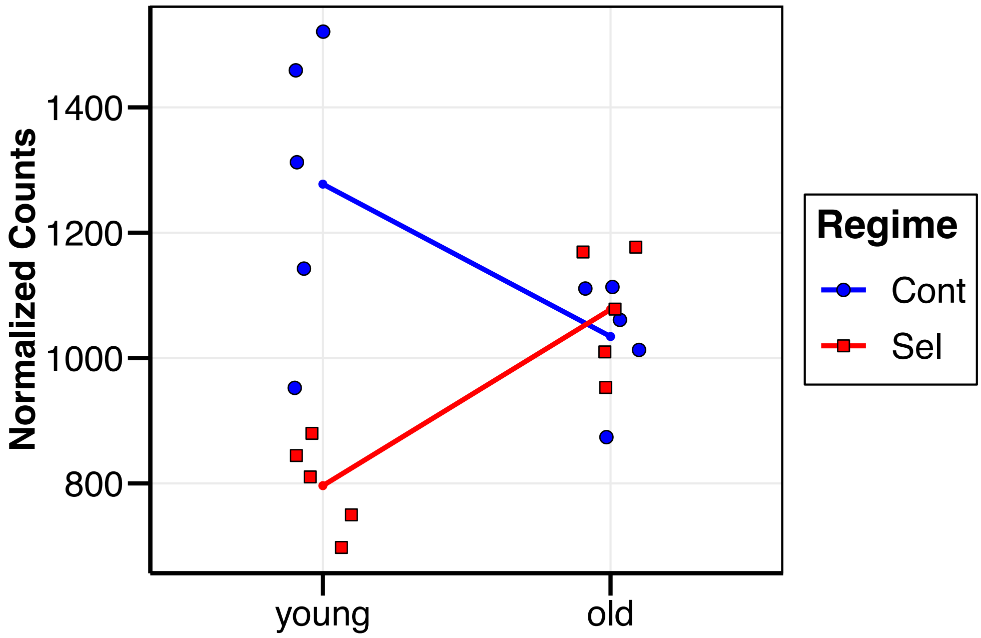

**Figure S14. Proportion of differentially expressed TEs and genes.** Distributions of differentially expressed TEs and genes across levels within Sex (M, males; F, females), Regime (S, selected; C, control populations), and Age (young; old) factors were compared using $\chi^2$ tests. TEs and genes were considered significant if their adjusted P-value was <0.05. n.s., not significant.

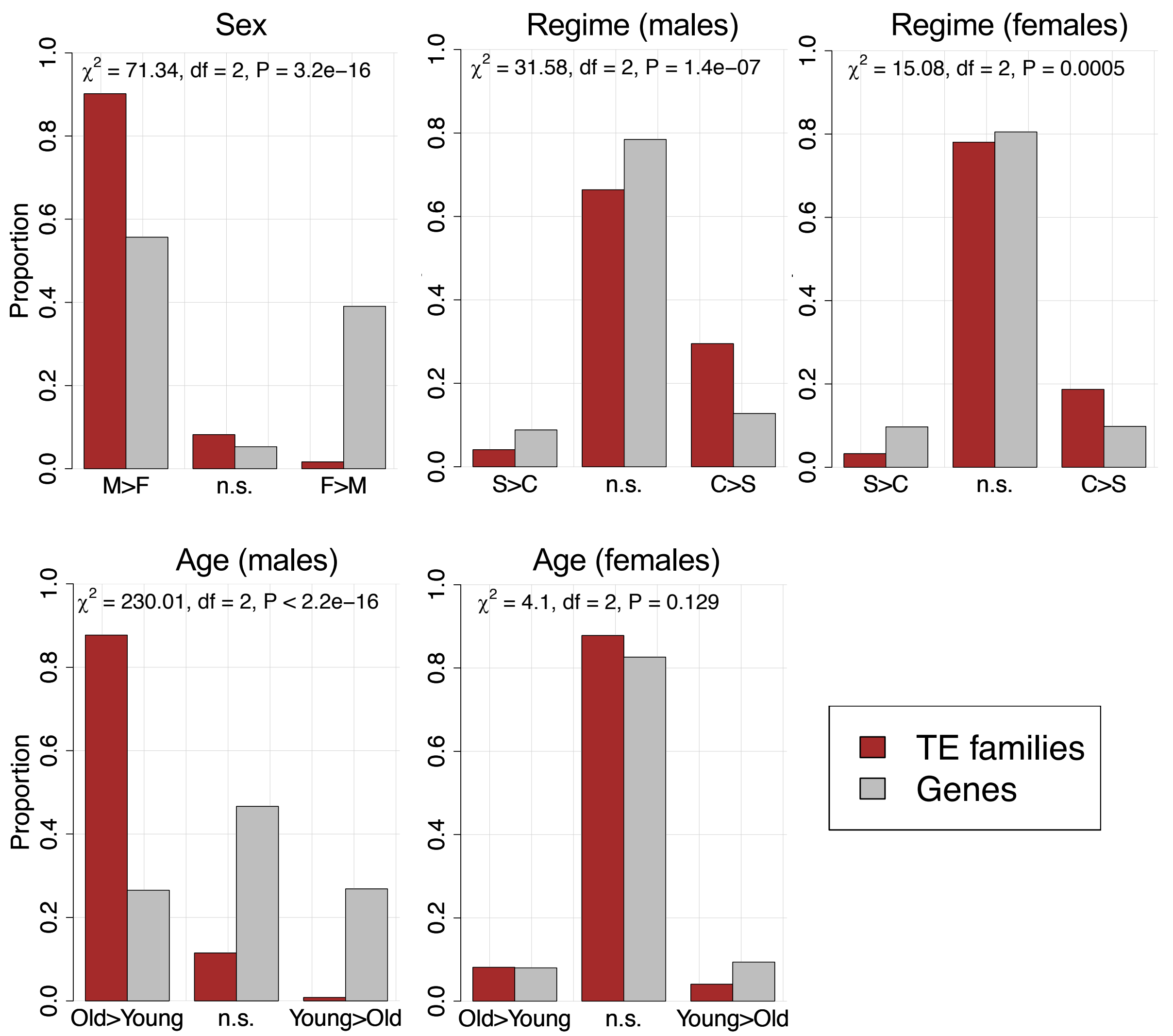

**Figure S15.** Normalized expression read counts obtained from DESeq2 were significantly correlated to the number of TE insertions in the genome. Expression values from females for each TE family were averaged across all conditions (age, regime). Separating samples into different conditions resulted in similar significant correlation coefficients (**Table S16**). ***Spearman's correlation ρ, P < 0.0001.

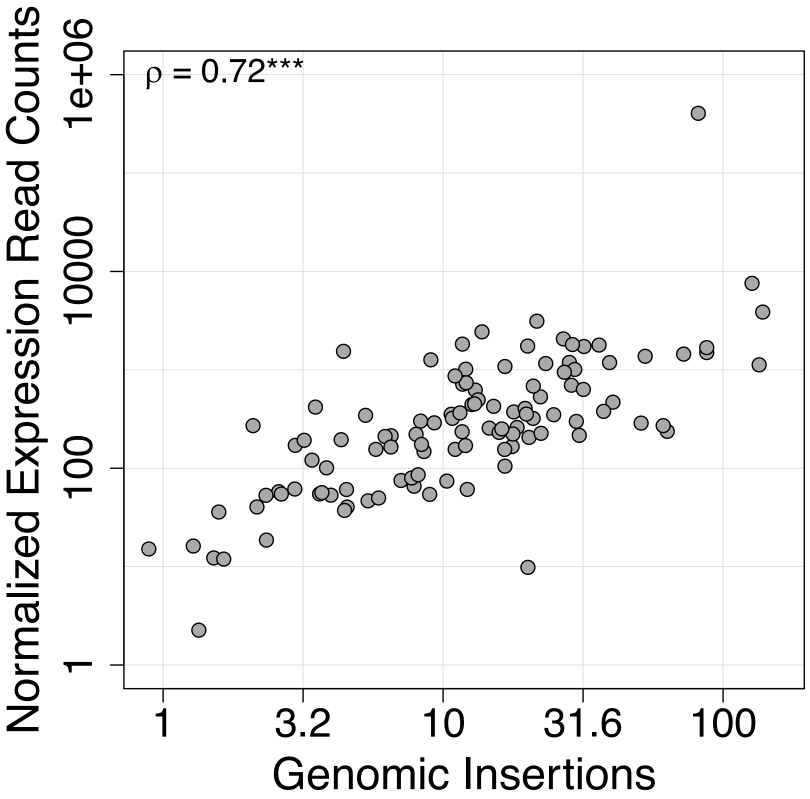
